# Physiological Harmony or Discord? Unveiling the Correspondence Between Subjective Arousal, Valence and Physiological Responses

**DOI:** 10.1101/2024.05.31.596899

**Authors:** Alina Koppold, Tina B. Lonsdorf, Manuel Kuhn, Mathias Weymar, Carlos Ventura-Bort

## Abstract

Affective experiences are inevitably accompanied by physiological changes, however it is still a matter of intense debate whether events evoking similar affective experiences produce comparable physiological responses (fingerprint hypothesis) or variation is the norm within individuals (population hypothesis). We reanalyzed data from two independent samples (N = 491; N = 64), using representational similarity analysis (RSA) to examine the trial-by-trial similarity patterns of subjective experience of valence and arousal and affect-related physiological measures (skin conductance [SCR] and startle blink responses). Across different affect-inducing (passive picture viewing, passive sound listening, imagery tasks) tasks and samples (Ns=491, 64), we observed strong-to-decisive evidence for a correspondence between SCR and startle responses and models of arousal and valence that assume variation, especially between trials generally evoking higher responses. Our results show that similar affective experiences are rather reflected by distinct physiological responses and emphasize the importance of considering intraindividual variability in future studies to better understand how physiological changes contribute to conscious affective experiences in humans.

---

Attending an electrifying concert, or watching a horror movie are experiences that evoke, almost ineluctably, affective subjective feelings accompanied by prominent bodily sensations (e.g., sweating, heart pounding). On a dimensional level, subjective affect (or affective state)— understood as the most basic consciously accessible feeling^1^— can be characterized by valence, referring to its hedonic value (ranging from pleasant to unpleasant), and arousal, referring to degree of activation (ranging from feeling calm to active, excited^1–7)^. While there is a general agreement on the characteristics of affect^1,6^, it is still unclear whether physiological changes mirror the experienced subjective affect. In other words, do similar affective experiences produce comparable physiological reactions within individuals? Here, we aimed at answering this fundamental question with potential implications in affective science^2^ and clinical fields^8–12^, by investigating the intraindividual correspondence between affective subjective experiences and physiological reactions in two independent, previously published^1^, samples^13,14^ across three well-known affect-inducing tasks^15^.

Previous research on the relationship between affective subjective experiences and physiological reactions has yielded conflicting views. Some suggest that experienced affect results from the activation of two motivational brain circuits—appetitive and defensive—that support survival^2,17–20^. The activation and intensity of these circuits are thought to determine experienced valence and arousal. When triggered (e.g., by life-relevant events), these circuits initiate autonomic changes that prepare the body for adaptive responses^20,21^. Consequently, physiological reactions should closely correspond with reported affect, leading to strong intraindividual consistency between experienced affect and physiological changes. Because this assumption aligns with the *fingerprint* hypothesis of discrete emotions^26,27^, which states that discrete emotions^2^ (e.g., happiness, sadness, fear) should have a prepared and distinctive physiological response^28^, we also label it the *fingerprint* hypothesis of affect (see Box).

Both in animals and humans, the skin conductance (SCR^29^) and the startle eye blink response^4,30,31^ have been pointed out as readout candidates of the activation of the appetitive and defensive motivational systems -and as correlates of the experienced arousal and valence, respectively. There is compelling evidence that SCR indicates the arousal level evoked by the triggering event^29,32–35^: high-arousing stimuli (both pleasant and unpleasant) tend to elicit stronger SCRs than their neutral counterparts. In contrast, the startle response is associated with valence, being potentiated for unpleasant stimuli and attenuated for pleasant ones, with higher arousal amplifying these effects^29,34,36–42^.

More recent views propose that although experienced affect and physiological changes are interconnected, they do not originate in the same underlying mechanism(s)^16,43–45,45^. These perspectives assume that autonomic reactions are strongly linked to the physiological needs of the current situation or context. Aligning with the principles of predictive coding^46,47^, they further suggest that physiological changes associated with an incoming event do not reflect an isolated response but are dependent on the interplay between the physiological demands of the current context and the predictiveness of the encountered event^61^. Importantly, these physiological reactions may partly subserve subjective affect. More concretely, experienced affect in response to a particular event may arise as the interaction between previous experiences with similar events, the ongoing read-out of the physiological bottom-up signals, and Bayesian inferences about the occurring event ^44, 66, 88, 75,89^. Crucially, due to the context and predictive heterogeneity of events categorized with similar affective attributes, the physiological needs in each of these instances may vary in a situated fashion. It is thus expected that within individuals, physiological variation is the norm across events evoking similar affective experiences. This view is in line with the *populations* hypothesis of discrete emotions^26,27^ (see also^51,52^) which states that emotions are not related to specific patterns of physiological activity. Thus, we refer to it as the *populations* hypothesis of affect (see Box). Recent reports have provided some evidence for within individual variability of physiological reactivity among events evoking similar affective experiences, supporting this hypothesis^26,14^.

Altogether, although prior studies pointed towards a strong correspondence between physiological reactivity and experienced affect^32,53^, supporting the *fingerprint* hypothesis of affect, emerging evidence suggests situated variability, in agreement with the *populations* hypothesis^26^. These somewhat opposing results may be related to different methodological factors. In many studies investigating a relationship between affective experience and physiological reactivity, findings are based on predefined categorical subdivisions (e.g., high vs low arousing; pleasant vs neutral vs unpleasant) across the dimensions of arousal and valence. This approach implies artificially transforming dimensional constructs into subcategories, which could overshadow the intertrial variability within categories^90^, and lead to ecological fallacy^54^, by assuming that the average value across trials holds for individual trials. Additionally, it prevents examining the association between affective experience and physiological reactions as a continuum, which becomes relevant to translate these findings to new understandings of psychological and clinical phenomena as dimensions^55^. Moreover, most of the studies have primarily focused on confirming the previously observed patterns rather than testing whether other models may explain the affect-physiology association more accurately^56^. The question thus arises as to whether these factors (i.e., disregarding intraindividual variability, lack of model testing) may pinpoint the correspondence between physiological reactions and affective experience in favor of one hypothesis over the other one.

Addressing these considerations is highly relevant, as it bridges clinical significance and scientific rigor. For instance, recent studies highlight the potential of intraindividual variability as a transdiagnostic feature of psychopathology. Individuals with post-traumatic stress disorder who have a history of childhood abuse exhibit heightened emotional variability, which correlates with both sudden gains in symptom improvement and therapeutic outcome^96^. However, greater variability is not universally beneficial. Higher negative affect variability has been repeatedly linked to maladaptive outcomes, including increased future negative affect^97^ anxiety disorders^98^, and greater severity of generalized anxiety disorder^99^. These findings suggest that intraindividual variability holds promise for advancing our understanding of psychopathology, particularly as a transdiagnostic marker, but also necessitates further investigation to uncover its underlying mechanisms.

Beyond its clinical relevance, intraindividual variability has implications for scientific rigor. The reliability paradox^100^ highlights that greater within-subject variability is associated with increased measurement precision but reduced statistical power^101^, pointing out the importance of intraindividual variability for assessing task replicability and generalizability.

A suitable tool to explore the complexities of intraindividual variability in the affect-physiology relationship is representational similarity analysis (RSA^57^). Instead of grouping trials in predefined categories, RSA allows for calculating the so-called representational (dis)similarity matrices (RSMs) which depict the relationship between trials on a dependent variable (e.g., SCR), dimensionally. Crucial to assess the role of intraindividual variability, RSMs can be calculated (1) taking into consideration inter-trial relationships within individuals, or (2) dismissing it (see Box).

Furthermore, RSA permits the creation of different distance models to test specific alternative hypotheses and determine which of them aligns with the data more closely^50^. Given these methodological advantages, here RSA was used to test for the similarity between physiological reactivity and subjective affect. Importantly, unlike previous studies, we directly compared the *fingerprints* and *populations* hypotheses of affect by creating models of subjective valence and arousal that partly align with each hypothesis. In particular, the Nearest Neighbors (NN) model was used to test for evidence in favor of the *fingerprint* hypothesis (similar affective experiences evoke similar physiological responses). The NN assumes that events evoking similar affective experiences produce comparable physiological responses. In addition, Anna Karenina (AK) models that predict variation within a dimension that align more with the *populations* hypothesis (similar affective experiences may evoke different physiological responses) were further used. These AK models, which were recently introduced to assume variation of individual differences in brain-behavior relationships^58^, predict similarities between dimensionally close trials on one side of a dimension, but dissimilarities on the other side of the dimension. More specifically, the AK model predicts similar physiological responding across high arousing/valence (positive) trials, but dissimilar physiological reactions across low arousing/valence (negative) trials, whereas the inverted AK model represents the opposite pattern (see Box).

In a first step, the role of within-individual variation on the affect-physiology correspondence was assessed by testing whether (1) considering vs (2) removing intraindividual variability would produce different RSMs. These RSMs were then compared with NN and AK-related models of subjective affective experience to test for the *fingerprint* and *populations* hypotheses, respectively.

If removing vs considering intraindividual variability leads to differential associations with the NN and AK-related models, the question arises as to which RSM (with or without intraindividual variability) is more representative of the individual data. That is, do individuals show more often one pattern than the other? The answer to this critical point would inform about which of the two RSMs more accurately represents the affect-physiology correspondence across participants. Hence, in a subsequent step, we tested the correspondence between the two RSMs (with or without individual variation) and the RSMs from each participant to elucidate which of the two patterns offers a more common representation of participants’ physiological responses.

## Results

### Skin Conductance Response (SCR)

To test the role of intraindividual variation on the affect-physiology correspondence, two averaged RSMs of SCR were created, one accounting for intraindividual (i.e., inter-trial) variability and the other dismissing intraindividual variability. Thereafter both averaged RSMs were compared to the averaged RSMs of arousal created based on NN and AK-related models. In the discovery sample (N= 483), which underwent a passive picture viewing task (PPV), both averaged RSMs of SCR (i.e., with and without intraindividual variability) were related to each other, rho = .08, p = .03, BF_10_ = 2.63, however, they showed differential relations with NN and inverted AK models of arousal.

### Averaged RSM of SCR disregarding intraindividual variability shows a strong correspondence with the averaged RSM of arousal based on the NN model

For the averaged RSM of SCR in which intraindividual variability was disregarded, decisive evidence was found for a positive association with the averaged RSM of arousal ratings based on the NN model, and anecdotal evidence for a lack of a relationship with the averaged RSM of arousal based on the inverted AK model (NN model: rho = .55, p <.001, BF_10_ > 100; inverted AK model: rho = −.06, p = .89, BF_10_ = 0.37). However, it should be noted that the NN and the inverted AK models make similar predictions in the lower end of the arousal dimension but opposite predictions in the upper end of the dimension. Indeed, both models are related to each other (rho = .25, p <.001, BF_10_ > 100). Hence, to test the correspondence with the NN and the inverted AK models after controlling for the shared similarities, we regressed one model from the other out and compared the correspondence between the model residuals and the averaged RSM of SCR. This led to decisive evidence for a correspondence between the averaged RSM of SCR dismissing intraindividual variability and the averaged RSM of arousal based on the NN model (rho = .58, p <.001, BF_10_ > 100), and decisive evidence for a lack thereof with the averaged RSM of arousal based on inverted AK model (rho = −.18, p =.999, BF_10_ =0.009; Figure 1A-C). These results indicate that, when no intraindividual variability is taken into account, the averaged pattern of the SCR is better explained by the NN model.

**Figure 1.**
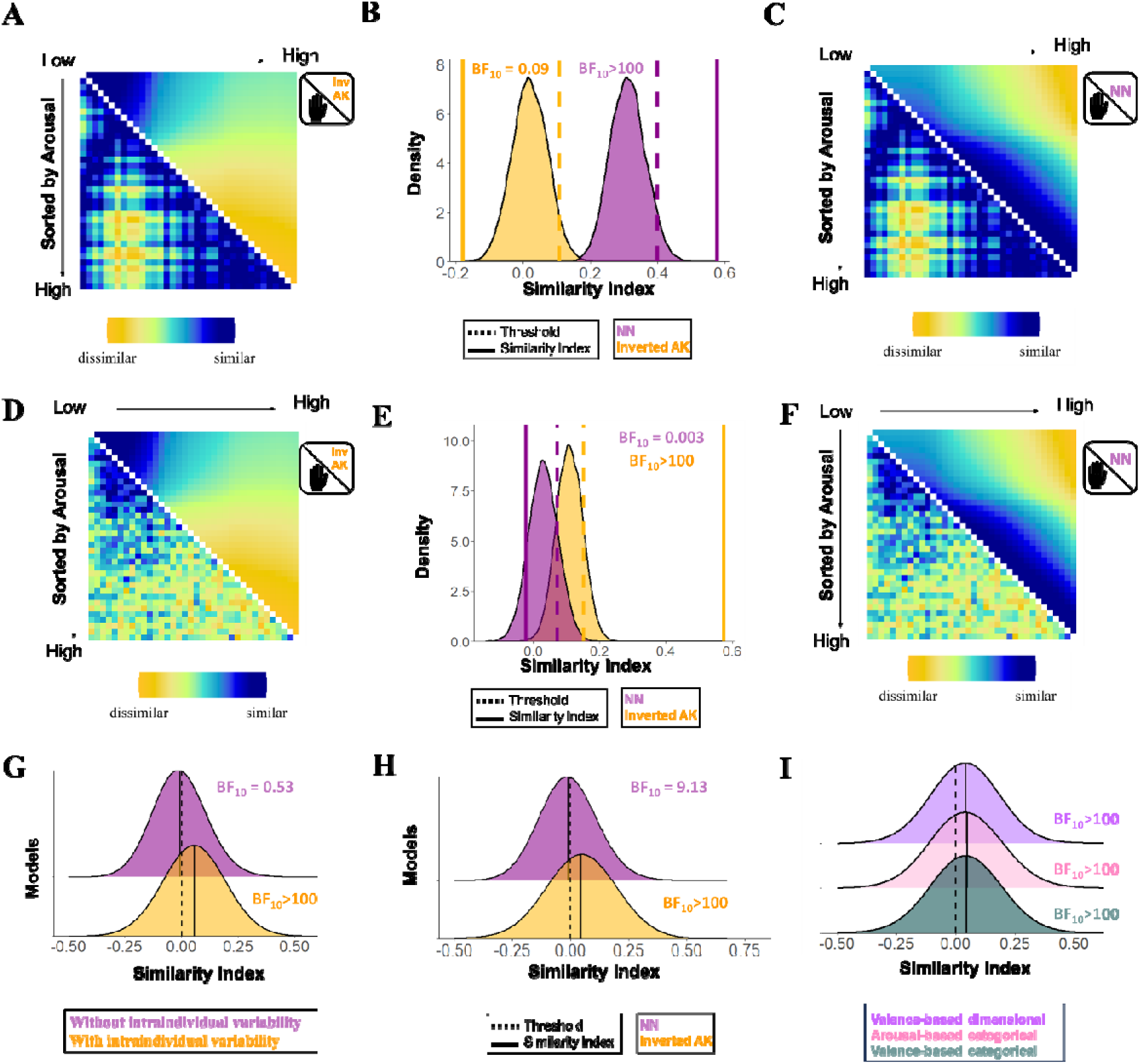
Association between the representational similarity matrices (RSMs) of subjective arousal based on the inverted Anna Karenina (inv AK) and Nearest Neighbors (NN) models (after controlling for the shared similarities), and the RSMs of the skin conductance response (SCR) dismissing (A-C) and considering (D-F) intraindividual variability (discovery sample, N= 483). A) Averaged RSM of SCR dismissing intraindividual variability (lower diagonal) and averaged RSM of subjective arousal based on the inverted AK model (upper diagonal). B) Results of the permutation test. C) Averaged RSM of SCR dismissing intraindividual variability (lower diagonal) and averaged RSM of subjective arousal based on the NN model (upper diagonal). D) Averaged RSM of SCR considering intraindividual variability (lower diagonal) and averaged RSM of subjective arousal based on the inverted AK model (upper diagonal). E) Results of the permutation test. F) Averaged RSM of SCR considering intraindividual variability (lower diagonal) and averaged RSM of subjective arousal based on NN model (upper diagonal). G) Distribution of the association between individual RSMs of SCR with the unique contribution of the averaged RSMs of SCR disregarding (purple) and considering (yellow) intraindividual variability. H) Distribution of the association between individual RSMs of SCR and unique contribution of the individual RSM of arousal based on the inverted AK and NN models. I) Distribution of the association between individual RSMs of SCR and individual RSMs of subjective arousal based on the inverted AK model after controlling for other models (i.e., valence-based categorical, arousal-based categorical, valence-based dimensional models).

### Averaged RSM of SCR considering intraindividual variability shows a strong correspondence with the averaged RSM of arousal based on the inverted AK model

For the averaged RSM of SCR considering intraindividual variability, we observed decisive evidence for correspondence with RSMs of both models (NN model: rho = .25, p <.001, BF_10_ > 100; inverted AK model: rho = .62, p <.001, BF_10_ >100), indicating that the averaged pattern of the SCR was similar to both the NN and the inverted AK models of arousal. Interestingly, when the similarity between models was regressed out, we observed decisive evidence for a correspondence between the averaged RSM of SCR and the averaged RSM based on inverted AK model (rho = .57, p <.001, BF_10_ >100), and decisive evidence for the lack thereof with the averaged RSM based on NN model (rho = −.02, p =.887, BF_10_ =0.003; Figure 1D-F). These results suggest that, when intraindividual variability was considered, the averaged pattern of the SCR was strongly related to the inverted AK model of arousal.

### Averaged RSM of SCR considering intraindividual variability provides a more accurate representation of individual physiological patterns

Given that both averaged RSMs of SCR (dismissing and considering intraindividual variability) showed differential associations with the NN and inverted AK models of arousal, in a follow-up analysis, we tested, which of the two averaged RSMs of SCR was a better representative of the individual RSMs of arousal. Results revealed evidence for a lack of a relationship between the individual RSMs of SCR and the averaged RSM of SCR dismissing intraindividual variability, Mean= .001, *t*(483)= 0.62, p = .53, BF_10_ = 0.53, however, decisive evidence was observed for a positive association between the individual RSMs and the RSM of SCR considering intraindividual variability, Mean = .054, *t*(483) = 12.34, p <.001, BF_10_ >100. These findings, which were further corroborated when models were regressed from each other out (RSM of SCR without intraindividual variability: Mean = −.01, *t*(483)= −2.12, p = .029, BF_10_ = 0.55; RSM of SCR with intraindividual variability: Mean = .055, *t*(483)= 12.34, p < .001, BF_10_ >100; Figure 1G), indicate that participants more often show a pattern of SCR similar to the averaged RSM of SCR that considers the individual fluctuations in physiological responses over trials.

### Individual RSMs of SCR show a strong correspondence with individual RSMs of arousal based on the inverted AK model

Additionally, individual level analysis was performed to test the relationship between individual RSMs of SCR and the individual RSM of subjective arousal based on the NN and the inverted AK models (see Box 1). Results showed evidence for a positive relationship of both the NN and the inverted AK models of subjective arousal with the individual RSMs of SCR (NN model: Mean = .014, *t*(482) = 3.93, p < .001, BF_10_ = 98.25; inverted AK model: Mean = .048, t(482)= 8.505, p <.001, BF_10_ >100). However, regression analysis revealed decisive evidence for a positive correspondence with the individual RSM of subjective arousal based on the inverted AK model (Mean = .047, *t*(482) = 8.72, p <.001, BF_10_ >100) and substantial evidence for a negative correspondence with the individual RSMs based on NN model (Mean = −.01, t(482)=-2.37, p = .018, BF10= 9.13; Figure 1H). Subsequently, we tested whether the evidence in favor of a relationship between the individual RSMs of SCR and the RMSs of subjective arousal based on the inverted AK model remained after controlling for other models, including valence- and arousal-based categorical, and valence-based dimensional models. In all cases, decisive evidence for a positive association between the inverted AK model and the SCR pattern was found after controlling for other models: relationship between SCR and inverted AK model of arousal after controlling for the valence-based categorical model, Mean = .044, t(482)= 8.08, p <.001, BF_10_ >100, the arousal-based categorical model, Mean = .046, t(482)= 8.23, p <.001, BF_10_ >100, and the valence-based dimensional model, Mean = .04, t(482)= 7.8, p <.001, BF_10_ >100; Figure 1I).

The replication sample (N= 64), which underwent a PPV, passive sound listening (PSL) task and Imagery task, allowed us to test whether the significant association between the RSMs of subjective arousal based on the inverted AK model and the RSMs of SCR would also be reproduced in another sample and could be generalizable across tasks. Results revealed a similar pattern of results to the discovery sample when participants were instructed to passively view affective images (PPV), listen to affect-inducing sounds (PSL) and to imagine affectively-laden events (Imagery task; Suppl. S4-S6), indicating that the correspondence between SCR and the inverted AK model of arousal was independent of the specific task and replicable across samples.

### Eye blink startle response

The two averaged RSMs of startle eye blink (dismissing and considering intraindividual variability) were compared with averaged RSMs of subjective valence based on models that align with the *fingerprints* (i.e., NN model) and with the *populations* hypothesis (AK-related models) of affect. In the discovery sample, both averaged RSMs of startle eye blink (i.e., with and without intraindividual variability) were related to each other, rho = .08, p = .38, BF_10_ = 0.29, and showed disparate relations with NN and AK models of valence.

### Averaged RSM of startle disregarding intraindividual variability shows a strong correspondence with the averaged RSM of valence based on the NN model

For the averaged RSM of startle blink responses in which intraindividual variability was disregarded, decisive evidence was found for a positive association with the averaged RSM of subjective valence based on the NN model, and anecdotal evidence for no relationship with the averaged RSM of valence based on the AK model (NN model: rho = .69, p <.001, BF_10_ > 100; AK model: rho = .05, p = .80, BF_10_ < .001). Subsequent regression analysis revealed decisive evidence for a correspondence between the averaged RSM of startle dismissing intraindividual variability and the averaged RSM of subjective valence based on the NN model (rho = .70, p <.001, BF_10_ > 100), and strong evidence for a lack thereof with the averaged RSM based on the AK model (rho = −.02, p =.998, BF_10_ =0.07; Figure 2A-C), indicating that, when no intraindividual variability was considered, the averaged pattern of the startle eye blink was better explained by the NN model.

**Figure 2.**
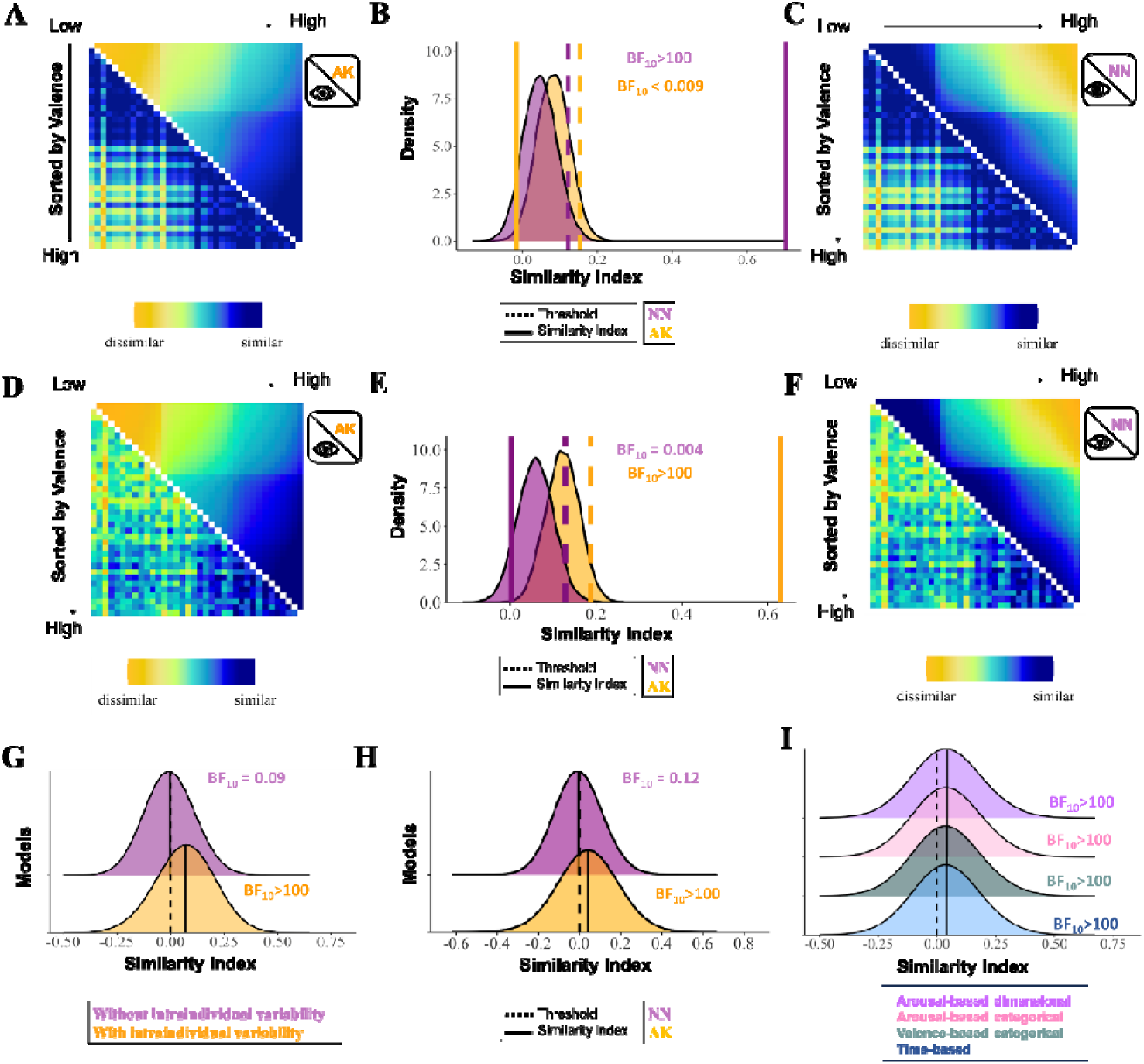
Association between the representational similarity matrices (RSMs) of subjective valence based on the Anna Karenina (AK) and Nearest Neighbors (NN) models (after controlling for the shared similarities), and the RSMs the startle eye blink response dismissing (A-C) and considering (D-F) intraindividual variability (discovery sample, N= 483). A) Averaged RSM of startle dismissing intraindividual variability (lower diagonal) and averaged RSM of subjective valence based on the AK model (upper diagonal). B) Results of the permutation test. C) Averaged RSM of startle dismissing intraindividual variability (lower diagonal) and averaged RSM of subjective valence based on the NN model (upper diagonal). D) Averaged RSM of startle considering intraindividual variability (lower diagonal) and averaged RSM of subjective valence based on the AK model (upper diagonal). E) Results of the permutation test. F) Averaged RSM of startle considering intraindividual variability (lower diagonal) and averaged RSM of subjective valence based on NN model (upper diagonal). G) Distribution of the association between individual RSMs of startle with the unique contribution of the averaged RSMs of startle disregarding (purple) and considering (yellow) intraindividual variability. H) Distribution of the association between individual RSMs of startle and unique contribution of the individual RSM of valence based on the AK and NN models. I) Distribution of the association between individual RSMs of startle and individual RSMs of subjective valence based on the AK model after controlling for other models (i.e., time-based, valence-based categorical, arousal-based categorical, valence-based dimensional models.

### Averaged RSM of startle considering intraindividual variability shows a strong correspondence with the averaged RSM of valence based on the AK model

When the averaged RSM of startle was computed considering intraindividual variability, an association was exclusively found with the averaged RSM of subjective valence based on the AK model (NN model: rho = .1, p = 0.32, BF_10_ = 2.03; AK model: rho = .62, p <.001, BF_10_ = 83.65; regression analysis: NN model: rho =0, p =.91, BF_10_ = 0.004 ; AK model: rho = .63, p <.001, BF_10_ >100; Figure 2D-F).

### Averaged RSM of startle considering intraindividual variability provides a more accurate representation of individual physiological patterns

Given the differential relationships between averaged RSMs of startle that dismiss vs consider intraindividual variability and the averaged RSM of subjective valence based on the NN and AK models we tested which of the two averaged RSM of startle was more closely related to the individual RSMs of startle. Results revealed no evidence for a positive relationship between the individual RSMs of startle and the averaged RSM of startle dismissing intraindividual variability, Mean = 0.007, *t*(398)= 2.5, p =.01, BF_10_ = 1.23, but decisive evidence for a positive relationship with the averaged RSM of startle that took into account intraindividual variability, Mean = 0.068, *t*(398)= 13.72, p <.001, BF_10_>100 (regression analysis: RSM of startle dismissing intraindividual variability: Mean = −0.003, *t*(398)= −.98, p = .32, BF_10_ = 0.09; RSM of startle considering intraindividual variability: Mean = 0.068, t(398)= 13.86, p <.001, BF_10_ >100; Figure 2G), indicating that participants more often show a startle pattern similar to the averaged RSM that takes into consideration intraindividual variability.

### Individual RSMs of startle show a strong correspondence with individual RSMs of valence based on the AK model

level analysis further demonstrated no association between individual RSMs of subjective valence based on the NN model and individuals RSMs of startle (Mean = .004, *t*(398)=1.47, p = .14, BF_10_ = 0.16) whereas individual RSMs of subjective valence based on the AK model were positively related to individual RSMs of startle (Mean = .041, *t*(398)=7.32, p <.001, BF_10_ >100). Regression analysis did confirm these findings (NN model: Mean = .002, t(400)=0.99, p = .32, BF_10_ = 0.12; AK model: Mean = .047 *t*(400)= 8.27, p <.001, BF_10_ >100; Figure 2H). Moreover, the association between individual RSMs of startle and individual RSM of subjective valence based on the AK model remained after controlling for other models, including time-based, valence- and arousal-based categorical, and valence-based dimensional models (time-based model: Mean = 0.042, t(400)= 7.63, p <.001, BF10 >100; valence-based categorical model: Mean = 0.041, t(400)=7.40, p <.001, BF10 >100; arousal-based categorical model: Mean = .043, t(400)= 7.65, p <.001, BF10>100; arousal-based dimensional model: Mean = .04, t(400)= 7.25, p <.001, BF10>100; Figure 2I).

In the replication sample, we could observe in the three tasks that the averaged RSM of startle considering intraindividual variability was a better representation of the individual RSM than the averaged RSM dismissing it. However, the associations with models of subjective valence were only partly reproduced in the PSL, but not in the PPV and Imagery tasks (Suppl. S7-S9).

## Discussion

It is widely accepted that affective experience could be conceptualized in terms of valence and arousal^1^. It is also suggested that the experience of affect is accompanied by a myriad of physiological changes. However, the correspondence between physiological responses to affective material and the experienced subjective valence and arousal is still a matter of intense debate. Whereas some evidence suggests that similar affective experiences generally evoke comparable physiological changes^20,25,32^ (*fingerprint* hypothesis of affect), there are also more recent studies that report within and between individual variability of physiological reactivity among events evoking similar affective experiences^14,26^ (*populations* hypothesis of affect). To advance the debate on the affect-physiology relationship, we used RSA to examine the trial-by-trial similarity patterns of the subjective affective experience (i.e., valence and arousal ratings) and physiological reactivity. SCR and startle blink responses were used as physiological correlates of arousal and valence, respectively. Capitalizing on the methodological features of the RSA, we compared the *fingerprints* and *populations* hypotheses of affect by creating models of valence and arousal that partly align with them (NN and AK/inverted AK, respectively). Furthermore, given that the role of intraindividual variability has often not been considered, we also investigated its influence on the relationship between physiological responses and experienced affect.

Regarding the arousal dimension, models that considered vs disregarded intraindividual variability in SCR were differentially associated with NN and inverted AK models of arousal in the discovery sample. When intraindividual variability was disregarded, we found decisive evidence for an association between the averaged RSM of SCR and the averaged RSM of arousal based on the NN model, whereas when intraindividual variability was considered, we observed decisive evidence for a relationship between the averaged RSM of SCR and the averaged RSM of arousal based on the inverted AK model. More importantly, follow-up analysis revealed that models of SCR that consider intraindividual variability represent the patterns of SCR across individuals more accurately than models dismissing it. Crucially, the results could be reproduced in all three affect-inducing tasks of the replication sample (Suppl. S4-S6), indicating that these findings are reliably observed regardless of the sample and task.

Results on the correspondence between the valence dimension and the startle eye blink response showed a similar pattern to arousal-SCR relationship. In the discovery sample, the averaged RSM of startle dismissing intraindividual variability was strongly associated with the averaged RSM of valence based on the NN model, whereas the averaged RSM of startle that considered intraindividual variability showed a strong correspondence with averaged RSM of valence based on the AK model. Subsequent analysis further revealed that the RSM of startle that considered intraindividual variability offered a more illustrative representation of individual startle response patterns across individuals. Although the association between models of valence and startle responses could not be confirmed in the replication sample, we observed in all three tasks that RSMs of startle that consider intraindividual variability offer a more representative depiction across individuals than RSMs that disregard it (Figures S6-S9 G).

These findings have two main implications for the relationship between physiological responses and experienced affect, in particular, and for the affective science field in general. First, they demonstrate the need to consider intraindividual variability across physiological responses to gain a more valid depiction of the physiological patterns that are evoked during affect-inducing tasks. To ensure that a specific physiological response (e.g., increase in SCR) is a readout measure of an ongoing psychological process (e.g., high levels of arousal) in a particular context (affect-inducing task), two important criteria need to be met^59^: (1) responses need to be specific, that is they should not occur if the psychological state does not take place and (2) they need to be reliable, that is they should often occur when the psychological state is given. If intraindividual variability is dismissed, the reliability and specificity of the responses remains unknown, which may lead to false conclusions and ecological fallacy^54,56,94^. This is particularly relevant in psychophysiological measures such as SCR, which has proven to have limited reliability at an individual level^95^. Here, we showed that disregarding vs considering this variability leads to somewhat different physiological patterns. Importantly, across physiological measures (SCR and startle), samples (discovery and replication samples), and tasks (PPV, PSL and Imagery tasks), we found compelling evidence for models that take intraindividual variability into account to depict a more exemplar (i.e., more often shared across participants) representation of the physiological patterns across participants. These findings provide empirical evidence that the timing of averaging within the analytical process is crucial in determining the extent of meaningful variance loss. This critical step should be meticulously reported in future studies ^94^. Our results further align with findings, showing the importance of person- and context-related variations to understand the relation between physiology and reported emotional experiences^26,56,60^.

Our results also display that dismissing vs considering intraindividual variability may lead to somewhat opposing interpretations in relation to experienced affect. When intraindividual variability was disregarded, the averaged RSMs of SCR and startle resembled models of arousal and valence that agree with the *fingerprint* hypothesis of affect whereas averaged RSMs that took into account intraindividual variability corresponded with models of arousal and valence that assume variation are in line with the *populations* hypothesis. This difference may help understand the disparity of results observed in the literature. During the last decades, the field of affective processing has been led by proposals that concur with the *fingerprints* hypothesis of affect^23^. These proposals assume that events that are rated similar in affective terms should produce similar physiological responses. Probably driven by this assumption, many of the previous studies on the affect-physiology relationship have opted for grouping responses from events that produced comparable affective feelings, not much focusing on intraindividual variability^29^. Consequently, many studies have shown that averaged physiological responses may covary with averaged affective reports, supporting the *fingerprint* hypothesis. However, more recent studies using data-driven multivariate methods that consider trial-by-trial relationships^14,26,56^ and thus account for intraindividual variability also indicate physiological variation among similar affective experiences, in line with the *populations* hypothesis.

The second major takeaway from our work is, that our findings demonstrate that the association between physiological responses and experienced affect is not harmonious, but partly discordant. The RSMs of physiology that provide a better depiction of the physiological patterns across participants revealed a similar pattern to models of affect that assume variation, in line with the *populations* hypothesis. More specifically, in the arousal dimension results showed that trials rated with low arousal levels, showing on average low physiological responsivity (i.e., lower or no SCR), produce comparable SCR. In contrast, in trials rated with high arousal levels, which are on averaged characterized increase in physiological responsivity, the evoked SCRs were more dissimilar to each other. An identical pattern was observed for the valence dimension, as trials higher in valence (associated with an averaged lower startle responsivity) showed comparable startle responses, while trials rated as lower in valence (i.e., more unpleasant) produced more dissimilar startle responses. These converging patterns indicate that the correspondence between physiological responses and subjective affective experience only emerge among trials generally evoking lower physiological responses, which may be due to floor effects (i.e., responses are too low to evidence differentiation). However, trials that when grouped are related to increase physiological responsivity, which are those high in arousal and low in valence for SCR and startle, respectively, are characterized by higher degrees of dissimilarity, indicating variation in the evoked physiological responses.

The present findings clearly reflecting physiological variability among trials with similar affective experiences (particularly high arousing trials for SCR and unpleasant trials for startle) are in line with theoretical views that concur with the principles of predictive processing^16,46,61–65^. These proposals postulate that physiological reactions associated with an incoming event do not reflect an isolated response, but depend on the physiological demands of the current situation and the interplay between the continuous physiological adjustments and the predictability of the encountered event^91,92^. These proposals further suggest that these physiological reactions are fed back to the brain by means of interoceptive mechanisms to keep the internal predictive models in check and update them, if necessary^46,61,66,67^. Importantly, these bottom-up signals (together with the myriad of sensory inputs) are incorporated as part of hierarchical, multidimensional abstractions to categorize the current state in affective terms. These proposals do not undermine the importance of physiological changes in the construction of conscious affective states but invite reframing the way the association between affect and physiological reactivity is currently outlined.

Previous studies have aimed at identifying physiological response patterns that could be indicative of a particular affective experience^4,29^ and whether variations in these physiological correlates could serve as individual predictors of psychopathological vulnerability^8,13,68,69^. Here, we demonstrated that events evoking similar affective experiences can produce dissimilar physiological responses (particularly in high arousing and low valence trials for SCR and startle, respectively), suggesting that physiological reactions do not accurately reflect an affective state (e.g.,^43^). Hence, rather than focusing on deciphering physiological patterns associated with specific affective states, future research may focus on understanding the mechanisms through which situated physiological changes are used to create conscious affective experiences and/or whether the disruption of these mechanisms may be a vulnerability factor for psychopathological disorders^70–72,96–99^, as recently demonstrated that affective disorders may be characterized by a divergence between autonomic responses and subjective emotional experiences^93^.

While our findings highlight the importance of considering intraindividual variability when assessing the relationship between physiological reactivity and affective experiences, shedding light on the discordant association between the experienced valence and arousal and physiological responses, it is important to acknowledge some limitations. In both the discovery and replication samples, the physiological reactivity and affective experience were not recorded at the same time (ratings were obtained separately after the presentation of the stimuli). Although affective experience to the same event may vary from moment to moment^73–75^, the fact that our pattern of results was consistently observed after controlling for other influential factors, including time (e.g., Figures 1I and 2I), suggests that this methodological caveat may not have a strong impact on the results. Nevertheless, future studies, using designs that simultaneously track affective experience and physiological reactivity^26,76,77^ may further provide valuable insights into the relationship between affective experience and physiological reactivity. The pattern of results observed for the SCR were replicated across samples and tasks but the association between startle and valence ratings based on the AK model found in the discovery sample was not replicated in the replication sample. The lack of replicability may be explained by differences in methodological characteristics. For instance, the discovery sample underwent only a single task, whereas the replication sample underwent three tasks, successively. Both studies also differed on the intensity of the startle probes (103 dB in discovery sample; 95 dB in replication sample). Hence, it could be speculated that longer durations of the experiment and/or lower startle probe intensities in the replication sample may have added variance to the physiological reactivity which may have shadowed the modulatory effects of valence (Suppl. S1). It is worth noting that while models used to test the *fingerprint* and *populations* hypotheses of affect align with some postulates of these hypotheses, they do not fully conform to them. For example, whereas the AK models predict variation on one side of the dimension, the *populations* hypothesis anticipates variation across the entire dimension. Despite these differences, we consider that the AK models remain valid for testing variation for the current psychophysiological measures given their unidirectional change. However, future studies employing a similar approach with other physiological variables that may show increasing and decreasing patterns (e.g., heart rate) may consider alternative models that predict variations throughout the dimension. Additionally, other models of affect beyond those here tested may also be considered^102^. In summary, this study validates two new RSA distance models (AK/ inverted AK) for peripheral psychophysiological data and highlights the necessity of considering intraindividual variability with multivariate approaches to create more representative physiological patterns across participant. This step is particularly crucial as we could demonstrate that considering vs. dismissing intraindividual variability may lead to somewhat opposing conclusions in the affect-physiology relationship. Furthermore, our results highlight a discordance between physiological responses and subjective affective experience, as suggested by the fact that events evoking similar affective experiences can produce dissimilar physiological responses. These findings can help advance the field in understanding how variations in physiological changes contribute to conscious affective experiences.

## Materials and Methods

### Participants

#### Discovery Sample

The sample comprises 685 individuals (424 female, 246 male; M_age_ = 25.36, SD_age_ = 5.57, from four sub-studies (for details^13^), which were conducted at the University Medical Center Hamburg-Eppendorf. Inclusion criteria encompassed physically and mentally healthy individuals between the ages of 18 and 50 who self-identified as of Caucasian origin. Exclusion criteria encompassed medication usage (except for oral contraceptives), a history of psychiatric diagnoses, and pregnancy. All participants had normal or corrected-to-normal vision and provided written informed consent in compliance with the protocol approved by the Ethics Committee of the General Medical Council Hamburg (PV 2755), adhering to the principles of the Declaration of Helsinki. Results on a different research question based on this dataset were published previously^13^.

#### Replication Sample

A total of sixty-four healthy students (51 women, 13 men; M_age_ = 23.43, SD_age_ = 4.11) from the University of Potsdam took part in the study in exchange of course credits or financial compensation. All participants had normal or corrected-to-normal vision. Each individual provided written informed consent for a protocol approved by the ethics committee of the University of Potsdam (28/2018). Before the experimental session, participants were screened and only invited to participate if they did not fulfil any of the following exclusion criteria: neurological or mental disorders, brain surgery, undergoing chronic or acute medication, history of migraine and/or epilepsy. The extended results from the replication sample were published somewhere else^14^.

### Material, design and procedure

#### Discovery Sample

The procedure of the passive picture viewing (PPV) task has been described previously elsewhere ^13,33^. In brief, the task consisted of two main phases: an initial 8-trial startle habituation phase (see below), followed by passive viewing of a set of 36 images. These images were selected from both the *International Affective Picture System* (IAPS)^15^ and *EmoPicS*^78^ databases, with an equal distribution of twelve images per emotional valence (negative, neutral, positive; detailed information can be found in^13,33^. The 36 images were presented twice, resulting in a total of 72 trials.

For positive valence, erotic images were tailored to match the subject’s sexual preference (i.e., same-sex or opposite-sex pictures), while negative images were selected to depict personally threatening situations (e.g., a gun pointed at the subject or an animal attacking the subject). Each image was displayed for 6 s, separated by randomized inter-trial intervals (ITIs) of 10, 12, or 14 s during which a fixation cross was presented. In 50% of the picture presentations, acoustic startle probes (103 dB, white noise) were presented either 4.5 or 5.5 s after the onset of the stimulus. To prevent predictability of the startle probes, 12 startle probes were distributed across the long ITIs (i.e., 14s), presented 8 s after ITIs onset.

During the initial startle habituation phase, eight startle probes (ITI: 11s) were presented while a fixation cross was displayed. Additionally, to maintain the subjects’ alertness, three occasional oddball trials were incorporated, as detailed in^33^. Post-experimentally, participants rated the previously seen pictures in terms of valence and arousal, using a computerized version of the self-assessment manikin (SAM^6^). Stimuli and SAM scales were presented simultaneously in the screen.

#### Replication Sample

The experimental session comprised five blocks. First, participants’ physiological responses were measured at rest for 6 minutes. The following three sections consisted of affect-inducing tasks, whose order was counterbalanced across participants, with different stimulus material: a passive picture viewing task (PPV), a passive sound listening task (PSL), and an imagery task. In the last block of the experimental session, participants rated the stimuli used in the three prior tasks in terms of valence and arousal, using a computerized version of the self-assessment manikin (SAM^6^).

#### Passive Picture Viewing (PPV) and Passive Sound Listening (PSL) tasks

A total of 36 images from the IAPS^15^ and 36 sounds from the *International Affective Digital Sounds* (IADS^79^) were selected based on their normative valence and arousal ratings and grouped in three affective categories (pleasant, unpleasant, and neutral). Each stimulus (image or sound presented for 6s) was followed by an ITI of 14, 16, or 18 s. In half of the trials, an acoustic startle probe (95 dB, white noise) was presented 4.5 or 5.5 s after cue onset. In half of the 18-second-long ITIs, the startle probe was also presented 9 s after cue offset (i.e., 18 startle probes during picture viewing/sound listening and 6 probes during ITIs). To account for startle habituation effects, two startle probes were presented one minute before each experimental block.

#### Imagery task

Eighteen scripts were selected from different sources: the *Affective Norms for English Text* (ANET; N = 15^15^), from prior studies^80^, and from in-house sets. All scripts were classified in three affective categories based on their normative valence and arousal ratings (for ANET scripts). Each script was presented twice, resulting in a total of 36 trials. Participants were instructed to read the scripts during presentation and, after script offset, to imagine themselves in the situation as vividly as possible. During the imagery period a circle was presented in the middle of the screen. As for the PPV and PSL tasks, the presentation order of the affective categories was pseudorandomized. The order of the scripts within each category was also pseudorandomized in 8 different orders. Scripts were presented for 12 s and followed by a 9-second imagery time window. The ITI varied between 14, 16, or 18 s. In 18 trials (once for each script), the acoustic startle probe was presented 4.5 or 5.5 s after onset of the imagery. Furthermore, six startle probes were presented 9 s after imagery cue offset in half of the 18-second-long ITIs. To account for startle habituation effects, two startle probes were presented one minute before the experimental block began.

#### Affective Rating task

The subjective valence and arousal ratings for each stimulus (36 images, 36 sounds or the 18 imagery scripts) were collected using the online platform *soscisurvey* (soscisurvey.de; ^54^). Which stimulus modality was rated first was dependent on the order of the tasks in the previous three blocks (e.g., if the participant underwent the PPV in block 2, the Imagery task in block 3 and the PSL in block 4, then images were rated first, then imagery scripts, and lastly sounds). Within each stimulus-set, the presentation of the stimuli was randomized. Participants were instructed to indicate on a SAM scale (1-9, see above) how they felt while viewing/listening/imagining the stimuli in terms of arousal and valence. Stimuli and SAM scales were presented simultaneously in the screen.

### Data recording, reduction and response definition

#### Discovery Sample

The recording of startle eyeblink responses and skin conductance responses (SCR) was conducted using a BIOPAC MP100 amplifier manufactured by BIOPAC Systems Inc. The data acquisition was managed using AcqKnowledge software version 3.9.2.

SCRs were assessed using self-adhesive Ag/AgCl electrodes, each measuring 55 millimeters in length. These electrodes were attached to the palmar side of the left hand (distal and proximal hypothenar regions). Prior to electrode placement, the hands were washed with warm water without the use of soap. SCRs were continuously recorded at a sampling rate of 1000 Hz and a gain of 5 micro-ohms (μΩ).

Subsequently, the recorded data were down-sampled offline to a rate of 10 Hz. SCR amplitudes, measured in microsiemens (μS), were then scored semi-automatically utilizing a custom-made program. The largest SCR response with an onset latency between 0.9 and 4.0 s after the onset of the stimulus was identified, as recommended by published guidelines^81^. To ensure normalization of the data, the SCR amplitude was log transformed (log[1+SCR amplitude]). All participants were included for the SCR analysis.

Fear-potentiated startle responses were recorded beneath the right eye through the use of two 4 mm Ag/AgCl electromyogram (EMG) electrodes strategically positioned over the orbicularis oculi muscle, along with one electrode placed on the participant’s forehead to serve as a reference point. The startle data were collected with a gain setting of 5000 at a rate of 1000 Hz. Online, the data underwent band-pass filtering within the range of 28 to 500 Hz, followed by rectification and integration (averaging over 20 samples).

The subsequent data analysis involved manual scoring based on a “foot to peak” approach, focusing on the time window from 20 to 150 ms following the onset of the startle probe. This scoring process adhered to established guidelines^82^ and was executed using the same program utilized for skin conductance responses (SCRs). Any responses affected by a blink occurring within 50 ms before the administration of the startle probe, recording anomalies, or excessive baseline activity (as determined by visual inspection) were categorized as missing values. All participants were included for the startle analysis.

Responses that were compromised by recording anomalies, such as electrode detachment or responses extending beyond the defined sampling window, were also excluded and considered as missing data points.

#### Replication Sample

##### Skin Conductance Response (SCR)

An MP-160 BIOPAC system (BIOPAC systems, Goleta, CA) with 2 shielded 8 mm Ag-AgCl sensors applied on the thenar and hypothenar eminence of the palmar surface of the participant’s left hand was used to collect electrodermal activity. SCRs were extracted using Continuous Decomposition Analysis in Ledalab Version 3.4.9^83^. The skin conductance data, initially recorded at a 2000 Hz sampling rate, were down sampled to a resolution of 50 Hz and parameters of the impulse response function (IRF) used to decompose the SC were optimized using four sets of initial values (see for details on Continuous Decomposition Analysis^83^). For the PPV and PSL task, the response window was set from 1 to 4 s after stimulus onset. For the Imagery task, the response window was set from 1 to 4 sec after the beginning of the imagery part. Both time-series and the averaged phasic driver^83^ were extracted as dependent variables. To ensure normalization of the data, the averaged phasic driver was log transformed (log[1+SCR driver]). Prior to performing the analysis, a visual inspection of the data was carried out to detect possible anomalies during recording. Due to anomalies during recording (excessive artifacts or near-to-zero skin conductance levels) 5 participants were excluded from the PPV task, 3 from the PSL task, and 3 from the Imagery task.

##### Startle Eyeblink Response

To measure the eyeblink component of the startle response, a MP-160 BIOPAC system (BIOPAC systems, Goleta, CA) was employed to record electromyographic (EMG) activity at sampling rate of 2000 Hz over the left orbicularis oculi, using two 4 mm Ag/AgCl miniature surface electrodes. To evoke the startle blink reflex, an acoustic startle probe, which consisted of 95 dB, 50 ms burst of white noise with a near instantaneous rise time, was used. The sound was administered in both ears using Audio Technica ATH-PRO700 MK2 headphones. Following EMG startle guidelines^82^, the data were filtered (high-pass filter of 60 Hz), rectified, and smoothed with a time constant of 10 ms. Startle blink responses were defined using in-house scripts in MATLAB 2020a. All trials were visually inspected and manually scored (c.f., guidelines for startle studies^82^). Startle responses were considered when starting 20-120 ms after probe onset and peaking within 150 ms. The magnitude of the eye blink response (in microvolts) was measured from onset to peak. Trials with no detectable blinks were scored as zero responses, and trials with excessive baseline activity (as determined by visual inspection) or recording artifacts were scored as missing. Participants with more than 50% of non-responses^55^ were excluded from the analysis (3 excluded participants from the PPV task, 3 from the PSL task, and 2 from the Imagery task).

### Representational similarity analysis (RSA)

#### Computation of Representational Similarity Matrices (RSMs) of models based on affective subjective experience

We defined three models of subjective experience of valence and arousal: the Nearest Neighbours (NN) model, the Anna Karenina (AK^58^) model and its inverted form (inverted AK). The NN assumes that a response on a specific trial is similar to responses on dimensionally close trials (e.g., low arousing trials should evoke similar responses) ^58^, implying that events evoking similar affective experiences produce comparable physiological responses in line with the *fingerprint* hypothesis. For instance, the NN model predicts similar SCRs among low arousing trials and among high arousing trials, separately, but dissimilar SCRs between high and low arousing trials. The AK-related models assume similarities between dimensionally close trials on one side of a dimension, but dissimilarities between dimensionally close trials on the other side of the dimension^58^. More concretely, the AK model represents higher similarity between trials located in the higher extreme of a dimension but higher dissimilarity between trials from the lower end of the dimension (e.g., similar physiological responding across high arousing/valence trials, but dissimilar physiological reactions across low arousing/valence trials). The inverted AK model represents the opposite pattern (e.g., similar physiological responding across low arousing/valence trials, but dissimilar physiological reactions across high arousing/valence trials). Given that the AK and inverted AK models are complimentary (they are perfectly, negatively related), to facilitate the interpretation of the results, we first visually inspected the physiological RSMs and determined the usage of the model which could resemble them most (i.e., showed more similarity rather than dissimilarity).

We constructed NN and AK-related models for arousal and valence ratings, using a two-step procedure. First, an RSM for each participant was constructed by sorting the trials based on the reported subjective experience (i.e., valence or arousal ratings, e.g., Jin et al., 2015) and calculating the trial-by-trial similarity scores. The formulae used for each model are as follows:

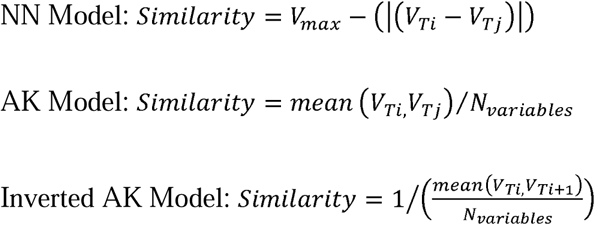

*V_max_* represents the maximum value of the dataset, and it is used as a constant to ensure that all similarity values stay in the positive range. *V_ti_* represents the score of trial *i. V_tj_*denotes the score of trial *j. N_variables_* indicates the number of trials (e.g., 36 or 18 trials).

In a second step, an averaged RSM was calculated by computing the average of the RSM across participants (so-called ‘averaged RSM’, see Box for terminology).

#### Computation of the physiological RSMs

##### SCR

As a dependent variable to calculate the RSMs, the SCR amplitude (log transformed to achieve normal distribution) was used in the discovery sample^3^ and the log transformed mean phasic driver (as an amplitude approximate derived from *Ledalab*^83^) in the replication sample. This approach for extracting the SCR may vary from our previous work, where the phasic driver vector was used (see Suppl. S2, S3 for additional comparative analyses).

##### Startle eyeblink response

To construct the RSMs of the startle, t-transformed magnitudes were used as dependent variables as suggested in the guidelines (Blumenthal et al., 2005). In trials in which startle response could not be quantified (i.e., missing trials), the similarity scores could not be calculated, and were thus treated as missing values. Although this strategy resulted in an uneven number of cases per participant, it prevented us from excluding participants with missing trials from the analysis.

To test for the effects of intraindividual variation, two different types of averaged RSMs of physiological responses were calculated, one disregarding intraindividual variability, and one considering it. To construct the averaged RSMs dismissing intraindividual variability, trials were first sorted based on the subjective affective ratings (SCRs were sorted based on subjective ratings of arousal and startle responses were sorted based on subjective ratings of valence) and trials were averaged across participants. The averaged RSM was calculated based on the resulting averaged scores for each trial (N trials = 36 for SCR in both samples, except for the Imagery task [N trials = 18]; N trials = 36 for the startle in the discovery sample and N trials = 18 in the replication sample).

For the construction of the RSMs of physiological responses considering intraindividual variability, we followed an identical procedure to the construction of the RSMs of the models of subjective affective experience. Trials were first sorted based on the individual subjective affective ratings of interest (i.e., arousal for SCR- and valence for startle-based RSMs) and the trial-by-trial similarity was calculated (i.e., using the NN formula) for each participant. This resulted in a single RSM for each individual participant. Thereafter, an averaged RSM was calculated by averaging the individual RSMs across the participants.

##### Computation of RSMs of alternative models

To ensure that the convergence between the RSMs based on subjective affective experience and the physiological RSMs was beyond other factors known to influence psychophysiological responses^84^ we computed RSMs based on the IAPS categorization of picture valence (pleasant vs unpleasant vs neutral) and arousal categories (low vs high arousing), and time (operationalized by the location in which trials position were presented). For the construction of the alternative RSMs, trials were initially sorted based on the subjective affective experience of interest (i.e., arousal for SCR- and valence for startle-based RSMs) and then the trial-by-trial similarities were calculated for each participant (using the NN-like formula), using as dependent variable nominal indexes for the category-based RSMs (e.g., for the valence category, pleasant images were coded as 1; unpleasant as 2; and neutral as 3; for the arousal category, high arousing were coded as 1 and low arousing as 2), and the trial number (e.g., from 1 to 36) for the time-based RSM.

##### Comparing model and physiological RSMs

Firstly, the two averaged physiological RSMs (without and with intraindividual variability) were compared to each other and to the averaged RSM of arousal and valence based on NN and AK models. More specifically, the averaged RSMs of subjective arousal based on the NN and inverted AK models were compared to the averaged RSMs of SCR, whereas the averaged RSMs of subjective valence based on NN and AK models were compared to the averaged RSMs of startle. The correspondence between averaged RSM of arousal and valence and the averaged RSMs of physiological responses was tested, using Spearman’s rank correlation coefficients, using the *stats* package in R. Because RSMs are mirrored along the diagonal, to avoid inflated correlations, only the values below the diagonal were considered for the correlation.

The statistical significance of the similarity between averaged RSMs of arousal and valence and the averaged RSMs of physiological responses was examined, using permutation tests by randomizing the modality labels of each pairing (i.e., model vs. physiological). The labels were iterated 10,000 times, using Monte Carlo random permutations. For each iteration, the correlation between RSMs was computed. These estimates served as a null distribution against which the actual correlation coefficient was compared. If the actual correlation coefficient was above 95% of the null distribution, we rejected the null hypothesis that the physiological and subjective RSMs are unrelated. We also used a Bayesian framework to quantify evidence in favor of the null (no positive relationship) or alternative (a positive relationship) hypotheses based on the results of the permutation test. To do so, we calculated the Bayes factor test for linear correlations, with the *BayesFactor*^85^ package in R environment (v. 4.3.3), using the distribution of the permutation test as the null interval.

Although the NN and AK and inverted AK models assume a different relationship between trials along a dimension (e.g., arousal), in some ranges of the dimension, they may show concurrent similarity patterns. For instance, for arousal the NN and inverted AK models show different patterns in high arousing trials, but similar patterns in low arousing trials. This implies that if a significant correspondence between the models and the physiological RSMs is observed, it may be unclear whether it is explained by the shared variance or by the uniqueness of each model. Thus, to further elucidate the unique contribution of each of the RSM models to the convergence with physiological RSMs, we performed regression analysis (e.g., the NN was regressed out from the inverted AK model and vice versa) and compared the correspondence between the residuals and the averaged physiological RSM^86^, using permutation tests and Bayesian testing.

To assess which of the averaged physiological RSMs (without and with intraindividual variability) was a better representative of the individual RSMs, in a subsequent step, individual RSMs were compared to both averaged RSMs (without and with intraindividual variability). To do so, similarity indexes (i.e., Spearman’s correlation coefficient) of the relationship between the individual RSMs and the averaged RSMs were extracted for each participant. The significance of the correspondence was tested by performing a one-sided t-test across participants, testing the null hypothesis of no correspondence (i.e., similarity does not differ from zero) between averaged and individual physiological RSMs. Bayesian testing was further performed to elucidate the evidence in favor of the null or alternative hypotheses. Given that both averaged RSMs may be related, we tested the unique contribution of each averaged RSM (i.e., without and with intraindividual variability). Similarity indexes of the relation between individual RSMs and the residuals of each of the averaged RSMs (extracted from regressing one model out of the other) were extracted for all participants and the correspondence was tested, using one-sided t-tests.

In addition, each participant’s physiological RSM was compared to the NN and AK-related models of arousal and valence. This was done by extracting similarity indexes between the physiological and subjective affective experience RSMs for each participant. The significance of the correspondence was tested by performing a one-sided t-test across participants, testing the null hypothesis of no correspondence (i.e., similarity does not differ from zero) between models and physiological RSMs. Bayesian testing was further performed to elucidate the evidence in favor of the null or alternative hypotheses. The unique contribution of the tested models (i.e., NN and inverted AK for SCR and NN and AK for startle) for the correspondence with physiological RSMs was further tested, using the abovementioned strategy on the model residuals (extracted from regressing one model out of the other). Additionally, we also tested whether the models explained variance beyond alternative models such as valence-based and arousal-based categorical models, time, and the alternative dimensional model (subjective valence for SCR and arousal valence for startle) by using the same regression procedure.

### Box Terminology

1. **The *fingerprints* hypothesis of affect** assumes a strong intraindividual correspondence between experienced affect and physiological changes. That is, in healthy individuals events evoking similar affective experiences are expected to produce similar physiological responses. The term *fingerprint hypothesis* was initially employed in the field of discrete emotions^26,27^ to group theories that assume that felt emotions should have consistent and specific physiological responses. Here, we decided to use the same term to refer to theoretical proposals advocating for consistent physiological reactions among events closed in the dimensions of valence and arousal.
2. **The *populations* hypothesis of affect** states that physiological variation is the norm across events evoking similar affective experiences. That is, events producing comparable affective experiences will evoke different physiological reactions. The term *populations hypothesis* was first used to group theoretical proposals stating that emotions, as abstract concepts formed by heterogeneous instances, are not related to specific patterns of physiological activity. Due to similarities with proposals assuming variation on the relation between physiology and affective subjective experience, we decided to use the same term to refer to the latter.
3. **Representational similarity matrices (RSMs) based on subjective affective experience** are created based on the valence and arousal ratings that participants provided for each event during the picture viewing, sound listening and imagery tasks. The formulae to calculate the similarities between trials depended on the model of interest: Nearest Neighbour (NN) model or Ana Karenina-(AK) related models).

a. ***NN Model*** assumes that a response on a specific trial is similar to responses on dimensionally close trials independently of their position on a dimensional scale (e.g., low, middle or high). We used the NN model to test for the *fingerprint* hypothesis of affect.
b. ***AK-related models*** assume that similarities between dimensionally close trials on one side of a dimension, but dissimilarities on the other side of the dimension. More concretely, the AK model represents higher similarity between trials located in the higher extreme of a dimension but higher dissimilarity between trials from the lower end of the dimension (e.g., similar physiological responding across high arousing/valence trials, but dissimilar physiological reactions across low arousing/valence trials), whereas the inverted AK model represents the opposite pattern. We used the AK-related models to test for the *populations* hypothesis.
4. **RSM based on physiological responses** are created based on the skin conductance (SCR) and startle eye blink responses. RSMs on physiological responses were calculated in two different ways:

a. ***Averaged RSM dismissing intraindividual variability.*** To calculate these RSMs, single trials are sorted by the variables of interest (e.g., arousal) and averaged across participants. Thereafter, the RSM of the averaged trials across participants is calculated.
b. ***Averaged RSM considering intraindividual variability.*** To create these RSMs, single trials are first sorted by the variable of interest (e.g., arousal), and then the RSMs of the physiological variable (e.g., SCR) calculated for each individual. Thereafter, individual RSMs are averaged together. This procedure was also used the create the averaged RSMs of subjective affect.
5. **Averaged level analysis** consists of the comparison between two averaged RSMs (e.g., averaged RSM of arousal based on the NN model and averaged RSM of SCR dismissing intraindividual variability). Averaged RSMs are compared, using Spearman’s correlation. Thereafter, this similarity index is compared to 10,000 random permutations (i.e., permutation test) to test for evidence for a positive association.
6. **Individual level analysis** is performed by first comparing two RSMs for each participant, using Spearman’s correlation (e.g., comparison between RSM of arousal based on the NN model and RSM of SCR for a specific subject), and then testing whether the similarity indexes are overall larger than 0, using simple t-tests.
7. **Regression analysis** is performed to test the relationship between two RSMs (e.g., averaged RSM of SCR dismissing intraindividual variability and averaged RSM of arousal based on NN model) after controlling for a third one (e.g., averaged RSM of arousal based on invAK model). RSMs (individual or averaged) are regressed out (e.g., the averaged RSM of arousal based on invAK is regressed out of the averaged RSM of arousal based on NN model) and the residuals are used to test for the similarity with the third RSM (e.g., the averaged RSM of SCR dismissing intraindividual variability) either using permutation tests (averaged level analysis) or testing whether the overall similarity indexes are larger than zero (individual level analysis).

## Supplementary material

### 1. Linear mixed models of the effects of Category on SCR and Startle

To ensure that the effects of the current data sets are in line with previous studies, showing affect-specific modulations in SCR and startle responses^1–3^, we carried out Linear Mixed Effects Models, using the lme4-package^4^. The categorical independent variable was valence category (i.e., pleasant, unpleasant or neutral; neutral was used as reference), whereas the dependent variable was z-transformed SCR and EMG amplitudes (discovery sample) or log-transformed mean phasic driver (replication sample) as well as t-transformed startle magnitudes (in both samples).

#### Discovery Sample

A significant main effect of picture valence emerged in both startle and SCR. For startle, the effect was significant (*F*(2, 20,077) = 268.48, *p*L<L.001), with a pattern of negative valence > neutral valence > positive valence (all pairwise comparisons *p*L<L.001). Similarly, for SCR, a significant effect was observed (*F*(2, 37,505) = 65.78, *p*L<L.001), with the valence gradient of negative > positive > neutral (all pairwise comparisons *p*L<L.002). startle responses aligned with the expected valence gradient, while SCR responses followed the expected arousal gradient.

#### Replication Sample

##### Skin Conductance Response

In the passive picture viewing (PPV) task, both pleasant, *t*(2,063)=7.25, *p* < .001, and unpleasant images, *t*(2,063)=6.42, *p* < .001, evoked larger SCRs compared to neutral images. In the passive sound listening (PSL) task, pleasant sounds evoked larger SCR than neutral ones, *t*(2,133)=4.32, *p* < .001, but no such effect was found for unpleasant relative to neutral sounds, *t*(2,133)=1.57, *p* = .12. In line with the PPV task, in the imagery task, both pleasant, *t*(2,133)=3.52, *p* < .001, and unpleasant scripts evoked larger SCR than neutral ones, *t*(2,133)=5.09, *p* < .001.

##### Startle eye blink response

In the PPV task, although pleasant and neutral pictures evoked comparable responses, *t(*1,088) =-0.75, *p* = .45, unpleasant images produced larger startle blink responses than neutral ones, *t(*1,088)=2.18, *p* = .029. In the PSL task, pleasant sounds evoked smaller responses than neutral ones, *t(*1,088)=-2.15, *p* = .032, but no significant difference was found for unpleasant relative to neutral sounds, *t*(1,088)=1.56, *p* = .117. In the Imagery task, unpleasant scripts evoked larger startle blink responses than neutral ones, *t*(1,107)=2.93, *p* = .003, but no differences emerged between pleasant and neutral scripts, *t*(1,107)=1.54, *p* = .12.

### 2. Comparison between time-series- and amplitude-based/phasic driver RSMs of SCR

In our previous study^5^, the RSMs of SCR were derived from the time-series of the SCR from 1 to 4 seconds after stimulus onset. This information was, however, unavailable in the discovery sample. Thus, in a preliminary step we tested the similarity between time-series- and amplitude-based and phasic driver RSMs of SCR in the replication sample. If both RSMs showed high correspondence, it would indicate that similar processes underlie both parameters, and their usage may be interchangeable. If that would not be the case, it would indicate that they are influenced by different factors. In the latter case, to determine which parameters to use, the consistency across tasks and samples was further tested (Suppl. S3).

For the PPV task, the similarity index (rho= −.04) did not surpass the permutation-based significant threshold (rho= .07, p = .786, BF_10_ = 0.001). A similar pattern was found in the PSL (similarity index: rho= −.02; threshold: rho= .07, p = .658, BF_10_ = 0.005) and Imagery tasks (similarity index: rho= −.18; threshold: rho= .14, p = .96, BF_10_ = 0.002). These results indicate that time-series and amplitude parameters of the SCR are influenced by different factors. Figure S1 depicts the RSM of time-series-(below the diagonal) and amplitude-based (above the diagonal) RSMs as well as the permutation test, comparing their correspondence in each of the three tasks.

**Figure S1.**
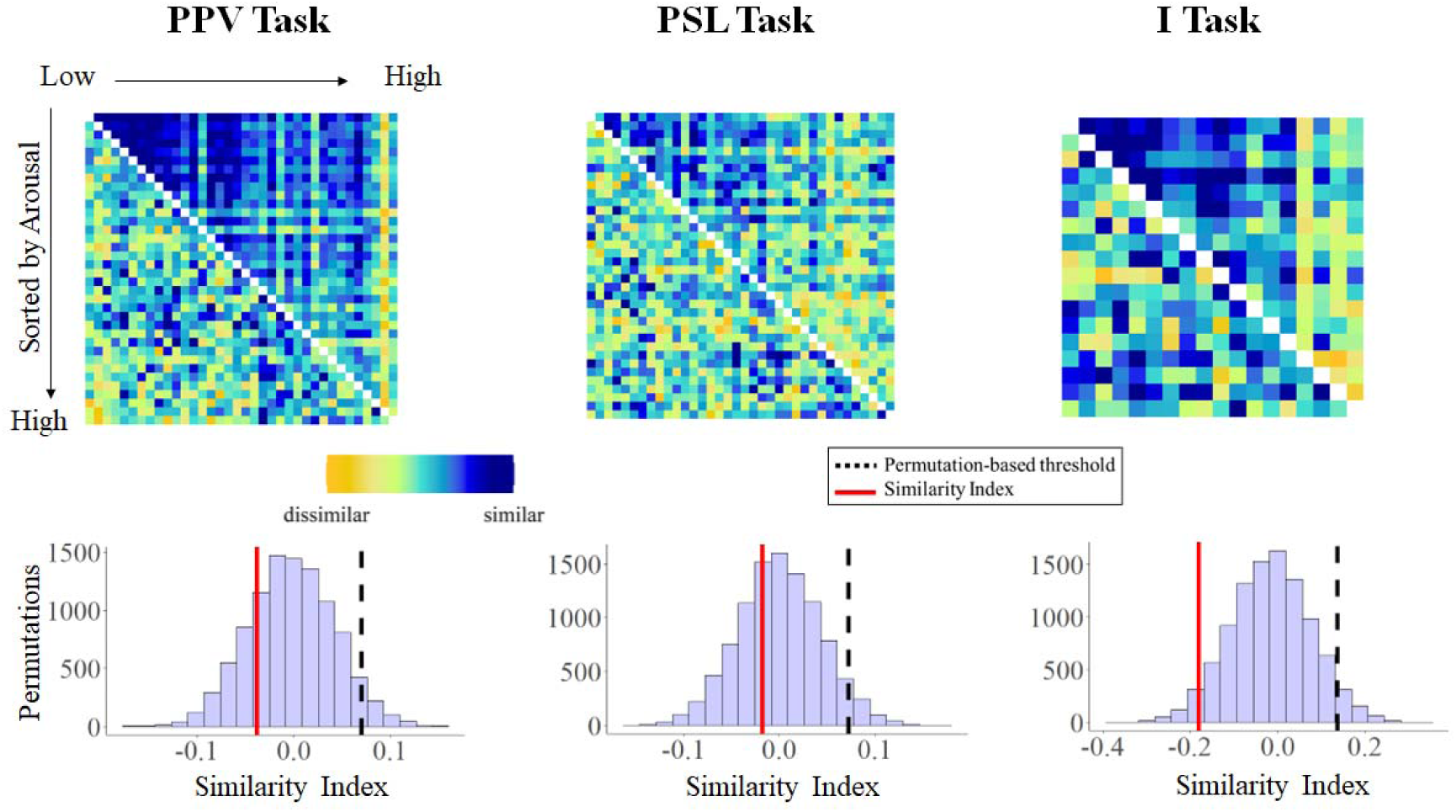
Results of the permutation tests comparing the RSMs derived from time-series and amplitudes of the SCR. In the upper part, the RSMs for both time-series (below the diagonal) and amplitudes (above the diagonal) are depicted for each task, PPV (right), PSL (middle), and Imagery tasks (left). In the lower part, the results of the permutation tests are depicted whereby the distribution of the permutation tests and the permutation-based threshold (black dotted line) as well as the similarity between both RSMs (red solid line) are represented. Results revealed no significant correspondence between RSMs across tasks, indicating that they are influenced by different factors.

### 3. Consistency of the time-series- and amplitude-based RSM of SCR across tasks and samples

Because no correspondence was found between time-series- and amplitude-based RSMs of SCR, in the following step, we examined the consistency of both parameters by comparing RSMs from the PPV and PSL tasks. For the time-series RSMs, results revealed no similarity between RSMs (rho= .01, threshold = .11, p = .84; BF_10_ = .002). For the amplitude RSMs, however, a significant correspondence was observed, rho = .18, threshold: rho = .11, p <.001, BF_10_ = 6.69). The amplitude-based RMSs not only showed consistency between tasks within the same sample, but also significant correspondence when comparing the RSMs of the PPV and PSL task with the PPV task of the discovery sample (PPV discovery sample and PPV replication sample: rho = .39, threshold: rho = .11, p <.001, BF_10_ > 100; PPV discovery sample and PSL replication sample: rho = .26, threshold: rho = .10, p <.001, BF_10_ > 100). Figure S2 depicts the RSM of the amplitude-based across tasks and samples as well as the permutation test, comparing their correspondence.

**Figure S2.**
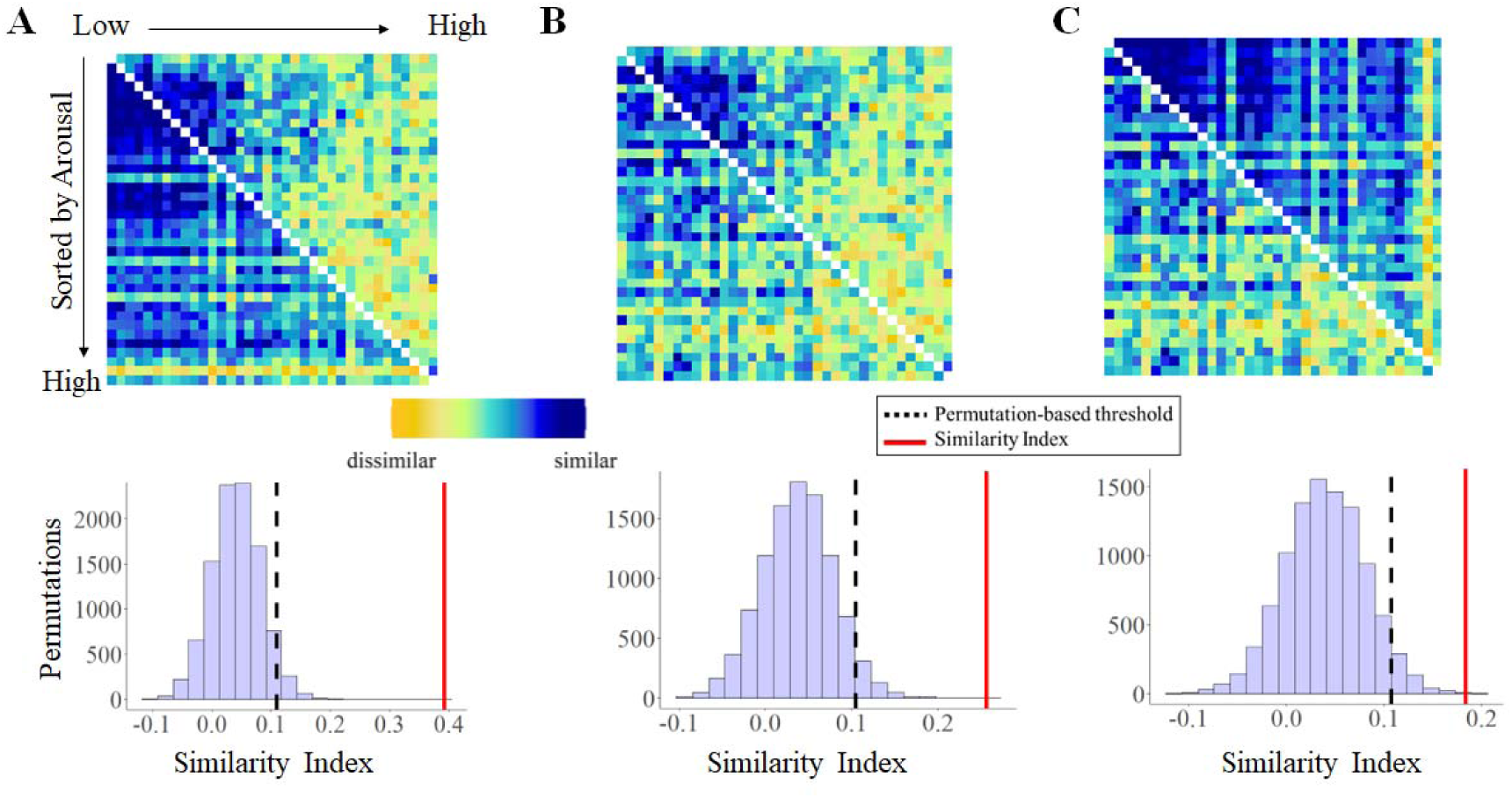
Results of the permutation tests comparing the RSMs derived from amplitudes of the SCR across tasks and samples. A) In the upper part, the RSMs for the SCR from the PPV of the discovery sample (above the diagonal) and from the PPV of the replication sample (below the diagonal). B) In the upper part, the RSMs for the SCR from the PPV of the discovery sample (above the diagonal) and from the PSL of the replication sample (below the diagonal). C) In the upper part, the RSMs for the SCR from the PPV (above the diagonal) and from the PSL of the replication sample (below the diagonal). In all sections (A, B, and C), the lower part depicts the results of the permutation tests whereby the distribution of the permutation tests and the permutation-based threshold (black dotted line) as well as the similarity between both RSMs (red solid line) are represented. Results revealed evidence for a correspondence between RSMs across tasks, and samples, indicating that they reflect similar processes.

### 4. Representational Similarity Analysis for the association between SCR and arousal-based models during the PPV task in the replication sample

In the PPV task of the replication sample (N = 64), a similar pattern of results was observed as in the discovery sample. Both averaged RSMs of SCR (i.e., considering and dismissing intraindividual variability) were related to each other, rho = .46, p <.001, BF10 >100, but showed different association with the NN and inverted AK models of arousal. For the averaged RSM of SCR dismissing intraindividual variability, decisive evidence for a positive association with the NN model of arousal was found, rho = .43, p <.001, BF_10_ >100, and strong evidence for a relationship with the inverted AK model of arousal, rho = .13, p = .015, BF_10_ = 71.27). Importantly, when models were regressed out, and the correspondence between the models residuals and the averaged RSM of SCR was tested, it was found decisive evidence for a correspondence between the averaged RSM of SCR dismissing intraindividual variability and the NN model (rho = .39, p <.001, BF_10_ >100), and moderate evidence for no correspondence with the inverted AK model (rho = −.01, p =.79, BF_10_ =0.14; Figure S3A-C). These results again indicate that when intraindividual variability is dismissed, the overall pattern of the SCR is better explained by the NN model.

When the averaged RSM of SCR considering intraindividual variability was compared to the models of arousal, a positive association with both the NN and inverted AK model was observed (NN model: rho = .10, p =.012, BF_10_ = 3.18; inverted AK model: rho = .59, p <.001, BF_10_ >100). However, regression analysis revealed decisive evidence for an association with the inverted AK model (rho = .55, p <.001, BF_10_ >100), and for a lack of a positive association with the NN model (rho = −.06, p = .998, BF_10_ = 0.001; Figure 2-E).

We further tested which of the averaged RSMs of SCR was a better representative of the individual RSMs of SCR. Results revealed decisive evidence for a positive relationship between the individual RSMs of SCR and the averaged RSM of SCR dismissing intraindividual variability, Mean = 0.09, t(58)= 6.07, p <.001, BF_10_ >100, and with the averaged RSM of SCR considering intraindividual variability, Mean = 0.17, t(58)= 9.67, p <.001, BF_10_ >100. Most importantly, when models were regressed out, the individual RSM were uniquely related to averaged RSM of SCR considering intraindividual variability (averaged RSM of SCR dis-missing intraindividual variability: Mean = −0.001, t(58)= −0.1, p = .92, BF_10_ = 0.14; averaged RSM of SCR considering intraindividual variability: Mean = 0.14, t(58)= 9.77, p <.001, BF_10_>100; Figure S3G), indicating that participants more often show a pattern of SCR similar to the averaged RSM of SCR considering intraindividual variability.

Individual level analysis revealed anecdotal evidence for a positive relationship between both the NN and the inverted AK and the individual RSM of SCR (NN model: Mean = .02, t(58)=2.06, p = .04, BF_10_ = 1.01; inverted AK model: Mean = .12, t(58)= 7.01, p <.001, BF_10_ >100) but regression analysis showed decisive evidence for the correspondence with the inverted AK model (Mean = .11, t(58)= 6.24, p <.001, BF_10_ >100) and no correspondence with the NN model (Mean = .008, t(58)= 0.69, p = .48, BF_10_ = 0.17; Figure S3H). The evidence for the association between the individual RSM of SCR and the inverted AK model remained after controlling for time, valence-based, and arousal-based categorical, as well as valence-based dimensional models (time-based model: Mean = .12, t(58)= 6.88, p <.001, BF_10_>100; valence-based categorical model: Mean = .08, t(58)= 7.00, p <.001, BF_10_ >100; arousal-based categorical model: Mean = .12, t(58)= 6.81, p <.001, BF_10_ >100; valence-based dimensional model: Mean = .11, t(58)= 6.9, p <.001, BF_10_ >100; Figure S3I).

**Figure S3.**
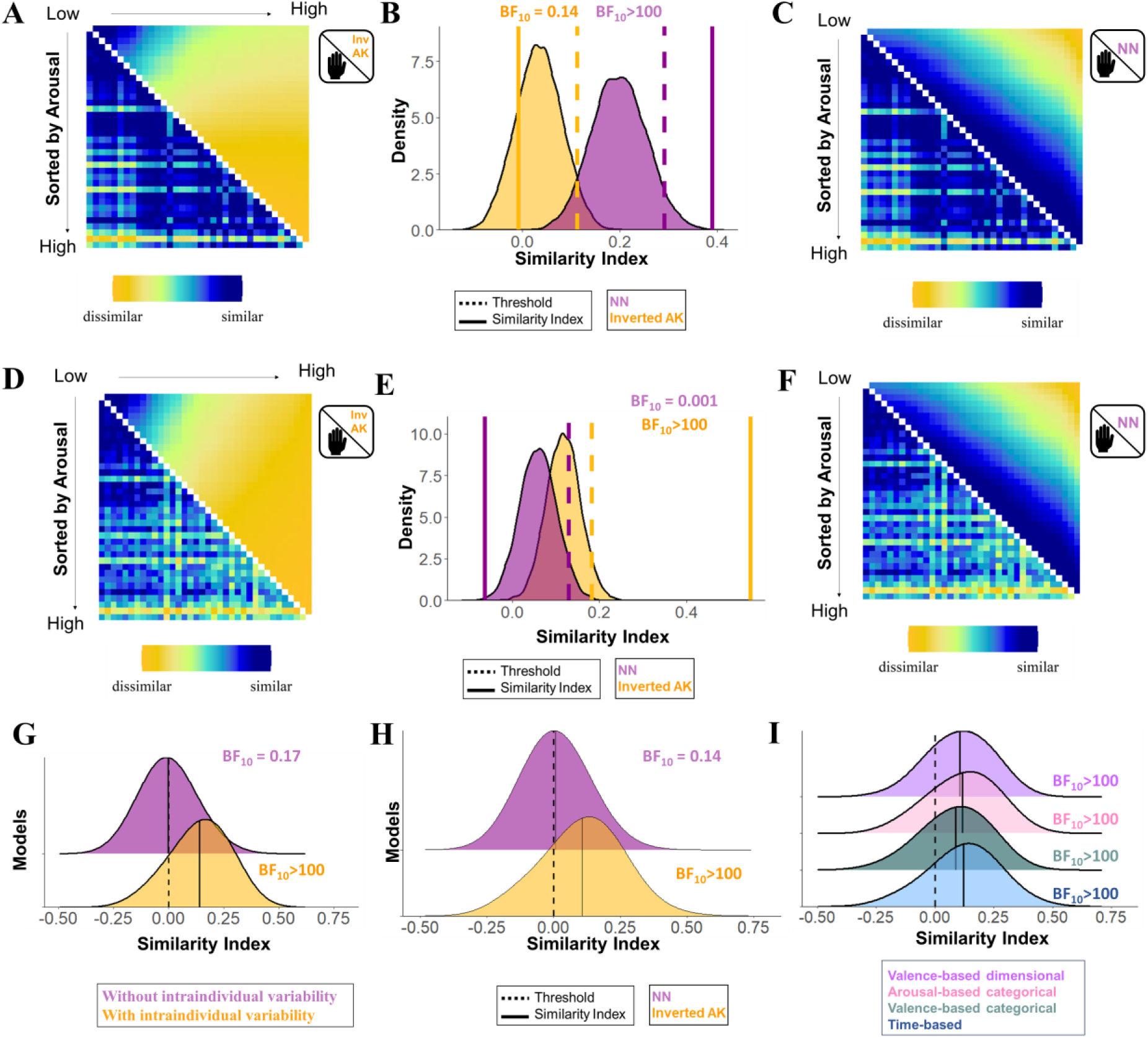
Association between the representational similarity matrices (RSMs) of subjective arousal based on the inverted Anna Karenina (inv AK) and Nearest Neighbors (NN) models (after controlling for the shared similarities), and the RSMs of the skin conductance response (SCR) dismissing (A-C) and considering (D-F) intraindividual variability (replication sample, N= 64, PPV task). A) Averaged RSM of SCR dismissing intraindividual variability (lower diagonal) and averaged RSM of subjective arousal based on the inverted AK model (upper diagonal). B) Results of the permutation test. C) Averaged RSM of SCR dismissing intraindividual variability (lower diagonal) and averaged RSM of subjective arousal based on the NN model (upper diagonal). D) Averaged RSM of SCR considering intraindividual variability (lower diagonal) and averaged RSM of subjective arousal based on the inverted AK model (upper diagonal). E) Results of the permutation test. F) Averaged RSM of SCR considering intraindividual variability (lower diagonal) and averaged RSM of subjective arousal based on NN model (upper diagonal). G) Distribution of the association between individual RSMs of SCR with the unique contribution of the averaged RSMs of SCR disregarding (purple) and considering (yellow) intraindividual variability. H) Distribution of the association between individual RSMs of SCR and unique contribution of the individual RSM of arousal based on the inverted AK and NN models. I) Distribution of the association between individual RSMs of SCR and individual RSMs of subjective arousal based on the inverted AK model after controlling for other models (i.e., time-based, valence-based categorical, arousal-based categorical, valence-based dimensional models).

### 5. Representational Similarity Analysis for the association between SCR and arousal-based models during the PSL task

In the PSL task of the replication sample (N = 64), both averaged RSMs of SCR (i.e., considering and dismissing intraindividual variability) were not related to each other, rho = .04, p =.13, BF_10_ = 0.43. For the averaged RSM of SCR dismissing intraindividual variability, decisive evidence for a positive association with the NN model of arousal was found, rho = .33, p <.001, BF_10_ >100, and decisive evidence for a lack of a positive relationship with the inverted AK model of arousal was observed, rho = −.14, p = .99, BF_10_ < 0.001). Importantly, when models were regressed out, and the correspondence between the model residuals and the averaged RSM of SCR was tested, strong evidence was found for a correspondence between the averaged RSM of SCR dismissing intraindividual variability and the NN model (rho = .36, p = .012, BF_10_ = 12.37), and decisive evidence for a lack thereof with the inverted AK model (rho = −.24, p =.99, BF_10_ < 0.001; Figure S4A-C). These results again indicate that when intraindividual variability is dismissed, the overall pattern of the SCR is better explained by the NN model.

When the averaged RSM of SCR considering intraindividual variability was compared to the models of arousal, moderate evidence for a lack of association between the averaged RSM of SCR and the NN model was found (rho = .06, p =.076, BF_10_ = 0.295) whereas decisive evidence for a correspondence with the inverted AK model was observed (rho = .45, p <.001, BF_10_ > 100). Regression analysis corroborated the exclusive correspondence with the inverted AK model (inverted AK model: rho = .45, p <.001, BF_10_ >100; NN model: rho = −.08, p = .993, BF_10_ = 0.004; Figure S4D-F).

We further tested which of the averaged RSMs of SCR (with or without considering intraindividual variability) was a better representative of the individual RSMs of arousal. Results revealed strong evidence for a lack of a positive relationship between the individual RSMs of SCR and the averaged RSM of SCR dismissing intraindividual variability, Mean = 0.003, t(60)= 0.35, p =.72, BF_10_ = 0.14, but decisive evidence was found for an association with the averaged RSM of SCR considering intraindividual variability, Mean = 0.12, t(60)= 8.9, p <.001, BF_10_ > 100. The results did not change when models were regressed out (averaged RSM of SCR dismissing intraindividual variability: Mean = −0.007, t(60)= −.88, p = .38, BF_10_ = 0.20; averaged RSM of SCR considering intraindividual variability: Mean = 0.12, t(60)= 9.03, p <.001, BF_10_ > 100; Figure S4G), indicating that participants more often show a pattern of SCR similar to the averaged RSM of SCR considering intraindividual variability.

Individual level analysis showed decisive evidence for a positive relationship between the inverted AK model and the individual RSM of SCR (Mean = .07, t(60)=4.01, p <.001, BF_10_ > 100), and evidence for the lack of a relationship with the NN model (Mean = .01, t(60)=1.25, p = .21, BF_10_ = 0.29). Further regression analysis validated these results (inverted AK model: Mean = .07, t(60)= 4.48, p <.001, BF_10_ > 100; NN model: Mean = −.007, t(60)= −0.65, p = .51, BF_10_ = 0.17, Figure S4H). The evidence for the association between the individual RSM of SCR and the inverted AK model was further observed after controlling for time, valence-based, and arousal-based categorical, as well as valence-based dimensional models (time-based model: Mean = .06, *t*(60)= 3.75, p <.001, BF_10_ > 100; valence-based categorical model: Mean = .07, *t*(60)= 4.22, p <.001, BF_10_ > 100; arousal-based categorical model: Mean = .06, *t*(60)= 3.82, p <.001, BF_10_ =76; valence-based dimensional model: Mean = .07, *t*(60)= 4.57, p <.001, BF_10_ >100; Figure S4I).

**Figure S4.**
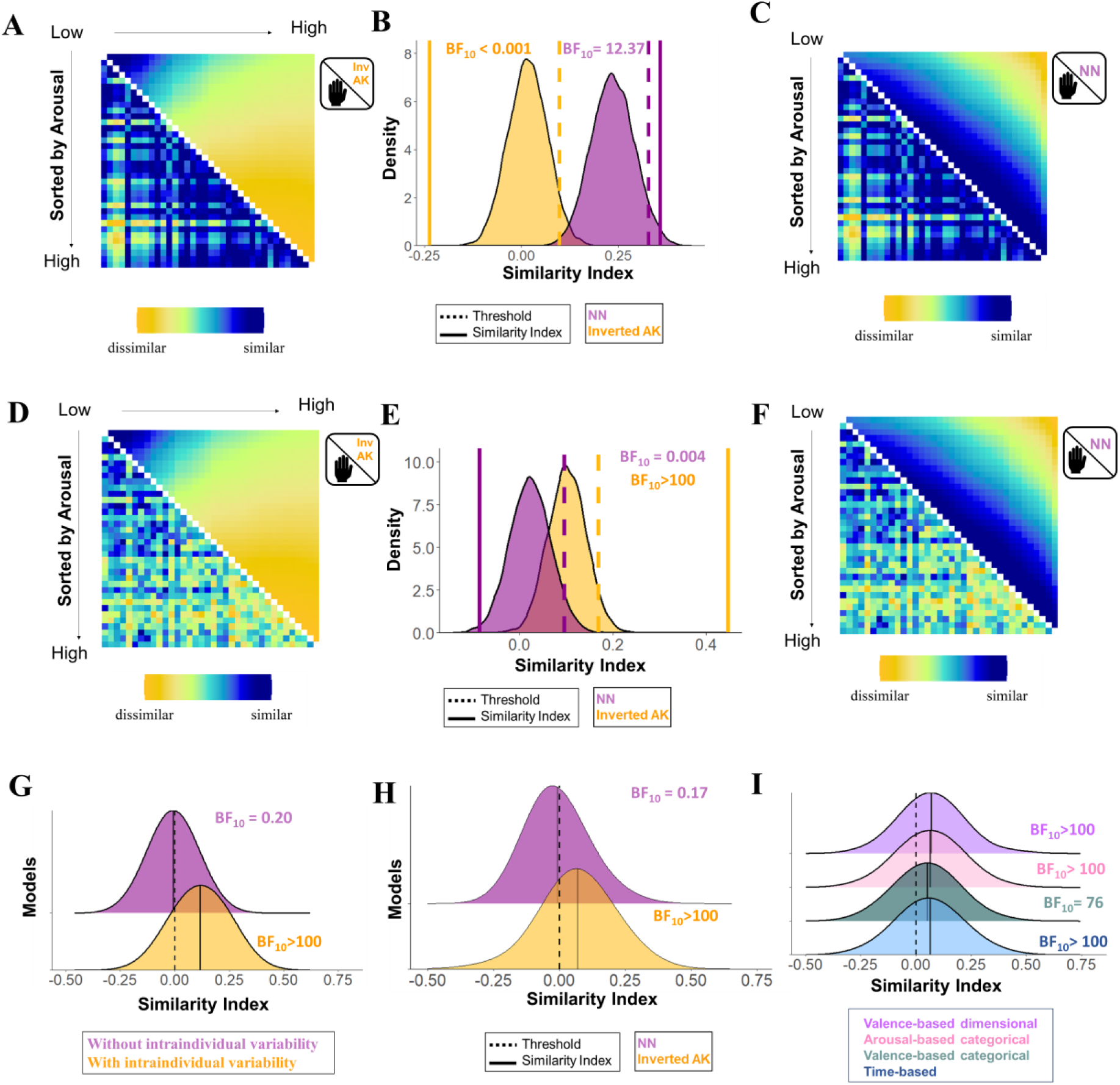
Association between the representational similarity matrices (RSMs) of subjective arousal based on the inverted Anna Karenina (inv AK) and Nearest Neighbors (NN) models (after controlling for the shared similarities), and the RSMs of the skin conductance response (SCR) dismissing (A-C) and considering (D-F) intraindividual variability (replication sample, N= 64, PSL task). A) Averaged RSM of SCR dismissing intraindividual variability (lower diagonal) and averaged RSM of subjective arousal based on the inverted AK model (upper diagonal). B) Results of the permutation test. C) Averaged RSM of SCR dismissing intraindividual variability (lower diagonal) and averaged RSM of subjective arousal based on the NN model (upper diagonal). D) Averaged RSM of SCR considering intraindividual variability (lower diagonal) and averaged RSM of subjective arousal based on the inverted AK model (upper diagonal). E) Results of the permutation test. F) Averaged RSM of SCR considering intraindividual variability (lower diagonal) and averaged RSM of subjective arousal based on NN model (upper diagonal). G) Distribution of the association between individual RSMs of SCR with the unique contribution of the averaged RSMs of SCR disregarding (purple) and considering (yellow) intraindividual variability. H) Distribution of the association between individual RSMs of SCR and unique contribution of the individual RSM of arousal based on the inverted AK and NN models. I) Distribution of the association between individual RSMs of SCR and individual RSMs of subjective arousal based on the inverted AK model after controlling for other models (i.e., time-based, valence-based categorical, arousal-based categorical, valence-based dimensional models).

### 6. Representational Similarity Analysis for the association between SCR and arousal-based models during the Imagery task

In the Imagery task of the replication sample (N = 64), both averaged RSMs of SCR (i.e., considering and dismissing intraindividual variability) were related to each other, rho = .47, p <.001, BF_10_ > 100. For the averaged RSM of SCR dismissing intraindividual variability decisive evidence for a positive association with the NN model of arousal was found, rho = .32, p <.001, BF_10_ >100, and moderate evidence for a positive relationship with the inverted AK model of arousal, rho = 0.21, p = .02, BF_10_ = 3.86). When models were regressed out, and the correspondence between the model residuals and the averaged RSM of SCR was tested, it was found moderate evidence for a lack of correspondence between the averaged RSM of SCR dismissing intraindividual variability and the NN model (rho = .28, p = .30, BF_10_ = 0.22), and anecdotal evidence for an association with the inverted AK model (rho = .16, p =.092, BF_10_ = 1.16; Figure S5A-C).

When the averaged RSM of SCR considering intraindividual variability was compared to the models of arousal, strong evidence for a lack of correspondence between the averaged RSM of SCR and the NN model of arousal was observed (rho = .01, p =.387, BF_10_ = 0.052), whereas decisive evidence for a correspondence with the inverted AK model of arousal was found (rho = .65, p <.001, BF_10_ > 100). Regression analysis corroborated these findings (NN model: rho = −.09, p = .998, BF_10_ = 0.042; inverted AK model: rho = .61, p <.001, BF_10_ >100; Figure S5D-F).

We further tested which of the averaged RSM of SCR was a better representative of the individual RSMs of SCR. Results revealed anecdotal evidence for a positive relationship between the individual RSMs of SCR and the averaged RSM of SCR dismissing intraindividual variability, Mean = 0.04, t(60)= 2.17, p =.03, BF_10_ = 1.25, but decisive evidence for an association with the averaged RSM of SCR considering intraindividual variability, Mean = 0.12, t(60)= 4.93, p <.001, BF_10_ >100. The results were even more pronounced when models were regressed out (averaged RSM of SCR dismissing intraindividual variability: Mean = −0.02, t(60)= −1.55, p = .11, BF_10_ = 0.46; averaged RSM of SCR considering intraindividual variability: Mean = 0.12, t(60)= 5.77, p <.001, BF_10_ >100; Figure S5G), indicating that participants more often show a pattern of SCR similar to the averaged RSM of SCR that considers intraindividual variability.

In line with the average analysis, in the individual level analysis decisive evidence in favor of a positive association between SCR and the inverted AK model (Mean = .10, t(60)=4.15, p <.001, BF_10_ > 100) and a lack thereof with the NN model was found (Mean = −.01, t(60)=-1.34, p = .18, BF_10_ = 0.33). Further regression analysis validated these results (NN model: Mean = −.02, t(60)= −1.91, p = .06, BF_10_ = 0.76, inverted AK model: Mean = .11, t(60)= 4.59, p <.001, BF_10_ > 100; see Figure S5D). The evidence for the association between the individual RSM of SCR and the inverted AK model was still observed after controlling for, valence-based, and arousal-based categorical, as well as valence-based dimensional models (valence-based categorical model: Mean = .10, t(60)= 4.25, p <.001, BF_10_ > 100; arousal-based categorical model: Mean = .1, t(60)= 4.13,, p <.001, BF_10_ > 100; valence-based dimensional model: Mean = .07, t(60)= 4.11, p <.001, BF_10_ >100).

**Figure S4.**
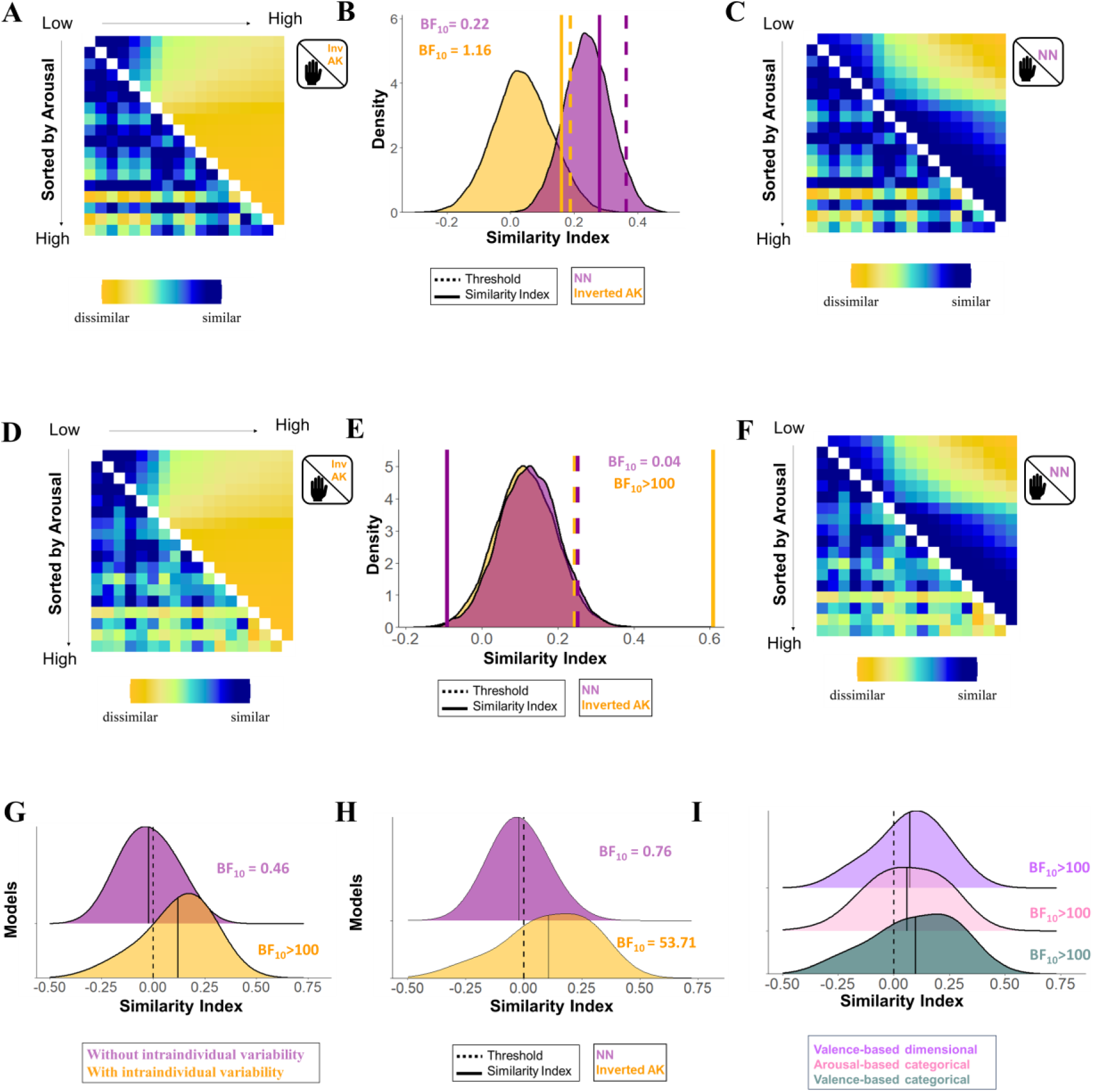
Association between the representational similarity matrices (RSMs) of subjective arousal based on the inverted Anna Karenina (inv AK) and Nearest Neighbours (NN) models (after controlling for the shared similarities), and the RSMs of the skin conductance response (SCR) dismissing (A-C) and considering (D-F) intraindividual variability (replication sample, Imagery task, N= 64). A) Averaged RSM of SCR dismissing intraindividual variability (lower diagonal) and averaged RSM of subjective arousal based on the inverted AK model (upper diagonal). B) Results of the permutation test. C) Averaged RSM of SCR dismissing intraindividual variability (lower diagonal) and averaged RSM of subjective arousal based on the NN model (upper diagonal). D) Averaged RSM of SCR considering intraindividual variability (lower diagonal) and averaged RSM of subjective arousal based on the inverted AK model (upper diagonal). E) Results of the permutation test. F) Averaged RSM of SCR considering intraindividual variability (lower diagonal) and averaged RSM of subjective arousal based on NN model (upper diagonal). G) Distribution of the association between individual RSMs of SCR with the unique contribution of the averaged RSMs of SCR disregarding (purple) and considering (yellow) intraindividual variability. H) Distribution of the association between individual RSMs of SCR and unique contribution of the individual RSM of arousal based on the inverted AK and NN models. I) Distribution of the association between individual RSMs of SCR and individual RSMs of subjective arousal based on the inverted AK model after controlling for other models (i.e., valence-based categorical, arousal-based categorical, valence-based dimensional models).

### 7. Representational Similarity Analysis for the association between startle and valence-based models during the PPV task

For the PPV task, both averaged RSMs of startle (i.e., considering and dismissing intraindividual variability) were related to each other, rho = .26, p <.001, BF_10_ = 15.64. For the averaged RSM of startle dismissing intraindividual variability, moderate evidence for a positive association with the NN model of valence was found, rho = .22, p = .009, BF_10_ = 4.61, and strong evidence for a lack of a positive relationship with the AK model of valence, rho = .09, p = .31, BF_10_ = 0.06. When models were regressed out, and the correspondence between the model residuals and the averaged RSM of startle was tested, it was found moderate evidence for correspondence between the averaged RSM of startle dismissing intraindividual variability and the NN model of valence (rho = .22, p = .007, BF_10_ = 5.56), and strong evidence for a lack of correspondence with the AK model of valence (rho =.09, p =.31, BF_10_ = 0.06; Figure S6A-C).

When the averaged RSM of startle considering intraindividual variability was compared to the models of valence, evidence for a lack of correspondence between the RSM of startle and the NN and AK models of valence was found at an average level (NN model: rho = .13, p = .08, BF_10_ = 0.83; AK model: rho = .10, p =.28, BF_10_ = 0.05; regression analysis: NN model: rho = .12, p = .08, BF_10_ = 1.01; AK model: rho = .10, p =.27, BF_10_ = 0.05; Figure S4D-F).

When the averaged RSMs of startle were compared to the individual RSMs, results revealed anecdotal evidence for a positive relationship between the individual RSMs of startle and the averaged RSM of startle dismissing intraindividual variability, Mean = 0.03, t(60)= 2.47 p =.01, BF_10_ = 2.31, but decisive evidence for an association with the averaged RSM of startle considering intraindividual variability, Mean = 0.11, t(60)= 8.38, p <.001, BF_10_ >100. The results were even more noticeable when models were regressed out (averaged RSM of startle dismissing intraindividual variability: Mean = 0.00, t(60)= 0.02, p = .98, BF_10_ = 0.14; averaged RSM of startle considering intraindividual variability: Mean = 0.11, t(60)= 7.58, p <.001, BF_10_ >100; Figure S6G), indicating that participants more often show a pattern of startle similar to the averaged RSM of startle considering intraindividual variability.

At an individual level, no evidence for a positive relationship between individual RSM of startle and models of valence was found (NN model: Mean = .01, t(60)=0.69, p = .49, BF_10_ = 0.16; AK model: Mean = .01, t(60)= 0.53, p =.59, BF_10_ =0.17; regression analysis: NN model: Mean = .02, t(60)=1.45, p = .15, BF_10_ = 0.38; AK model: Mean = .02, t(60)= 0.99, p =.32, BF10=0.22; Figure S6I). Moreover, similar evidence for a lack of association between the individual RSM of startle and the AK model was found after controlling for time-based (Mean = .01, t(60)= 0.31, p = .75, BF_10_ =0.14), valence-based categorical (Mean = .01, t(60)= 0.48, p =.63, BF_10_ = 0.16), arousal-based categorical models (Mean = .01, t(60)= 0.49, p =.62, BF_10_ = 0.16) and arousal-based dimensional model (Mean = .01, t(60)= 0.43, p = .67, BF_10_ = 0.15).

**Figure S6.**
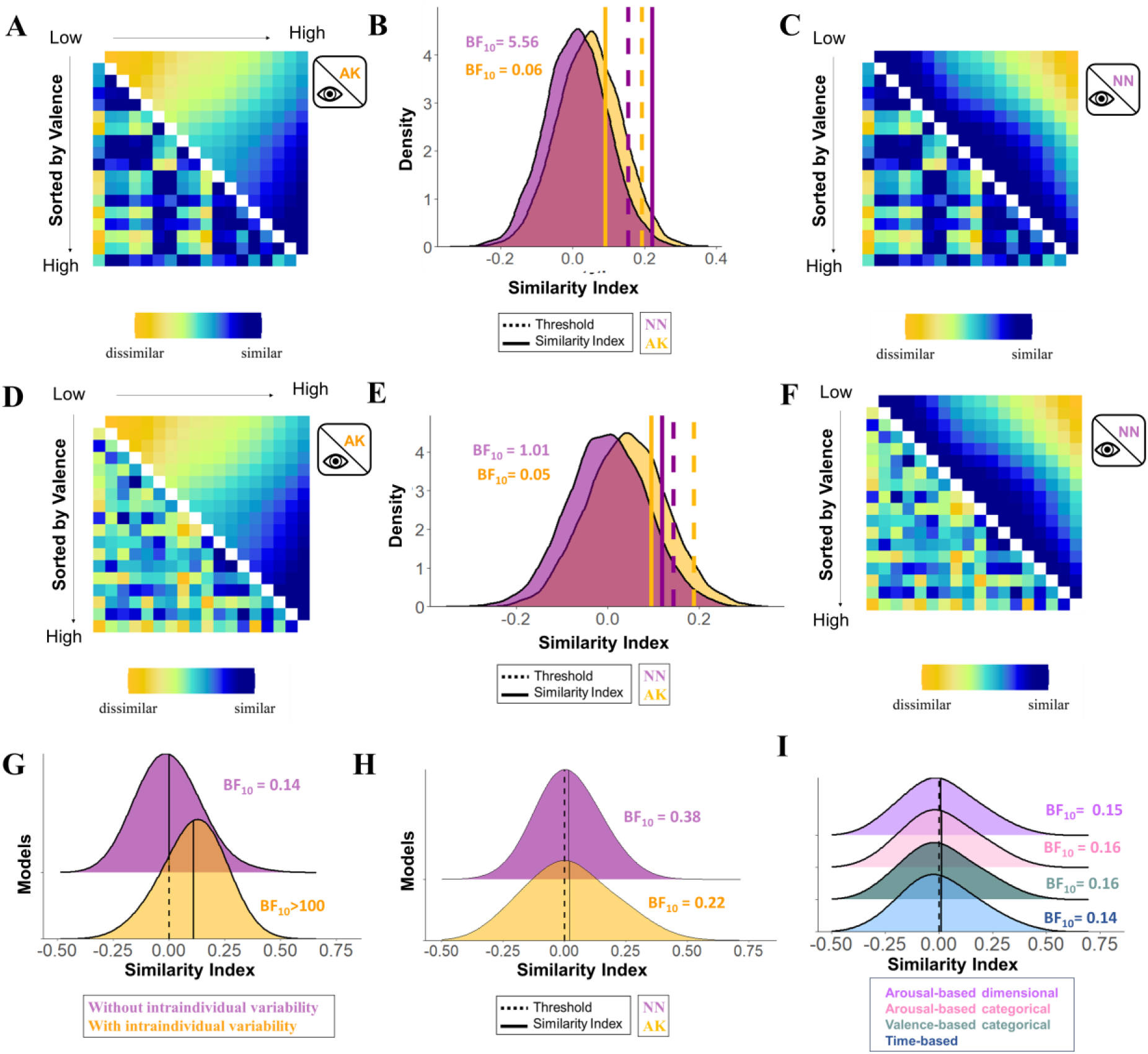
Association between the representational similarity matrices (RSMs) of subjective valence based on the Anna Karenina (AK) and Nearest Neighbours (NN) models (after controlling for the shared similarities), and the RSMs the startle eye blink response dismissing (A-C) and considering (D-F) intraindividual variability (replication sample, PPV task, N= 64). A) Averaged RSM of startle dismissing intraindividual variability (lower diagonal) and averaged RSM of subjective valence based on the AK model (upper diagonal). B) Results of the permutation test. C) Averaged RSM of startle dismissing intraindividual variability (lower diagonal) and averaged RSM of subjective valence based on the NN model (upper diagonal). D) Averaged RSM of startle considering intraindividual variability (lower diagonal) and averaged RSM of subjective valence based on the AK model (upper diagonal). E) Results of the permutation test. F) Averaged RSM of startle considering intraindividual variability (lower diagonal) and averaged RSM of subjective valence based on NN model (upper diagonal). G) Distribution of the association between individual RSMs of startle with the unique contribution of the averaged RSMs of startle disregarding (purple) and considering (yellow) intraindividual variability. H) Distribution of the association between individual RSMs of startle and unique contribution of the individual RSM of valence based on the AK and NN models. I) Distribution of the association between individual RSMs of startle and individual RSMs of subjective valence based on the AK model after controlling for other models (i.e., time-based, valence-based categorical, arousal-based categorical, valence-based dimensional models).

### 8. Representational Similarity Analysis for the association between startle and valence-based models during the PSL task

For the PSL task of the replication sample (N = 64), both averaged RSMs of startle (i.e., considering and dismissing intraindividual variability) were related to each other, rho = .3, p <.001, BF_10_ = 12.39. For the averaged RSM of startle dismissing intraindividual variability, anecdotal evidence for a lack of a positive association with the NN model of valence was found, rho = .14, p = .063, BF_10_= 0.80, and strong evidence for lack of a positive relationship with the AK model of valence, rho = −.001, p = .65, BF_10_= 0.02. When models were regressed out, and the correspondence between the model residuals and the averaged RSM of startle was tested, it was found anecdotal evidence for a lack of correspondence between the averaged RSM of startle dismissing intraindividual variability and the NN model of valence (rho = .14, p = .16, BF_10_= 0.93), and strong evidence for a lack of correspondence the AK model of valence (rho =-.002, p =.67, BF_10_= 0.02; Figure S7A-C).

When the averaged RSM of startle considering intraindividual variability was compared to the models of valence, anecdotal evidence for a positive association between the AK, but not the NN model, and the averaged RSM of startle was found (AK model: rho = .23, p =.018, BF_10_= 1.53; NN model: rho = −.09, p = .874, BF_10_= 0.01). These results were further verified in the regression analysis (NN model: rho = −.10, p =.887, BF10=0.002; AK model: rho = .23, p =.018, BF_10_= 1.68; Figure S7A-C).

We also tested which of the averaged RSM of startle was a better representative of the individual RSMs of startle. Results revealed anecdotal evidence for a positive relationship between the individual RSMs of startle and the averaged RSM of startle dismissing intraindividual variability, Mean = 0.03, t(60)= 2.21 p =.03, BF10 = 1.35, but decisive evidence for an association with the averaged RSM of startle that considering intraindividual variability, Mean = 0.12, t(60)= 7.91, p <.001, BF_10_>100. The results were even more pronounced when models were regressed out (averaged RSM of startle dismissing intraindividual variability: Mean = −0.003, t(60)= −0.19, p = .84, BF_10_= 0.84; averaged RSM of startle considering intraindividual variability: Mean = 0.12, t(60)= 7.52, p <.001, BF_10_>100; Figure S7G), indicating that participants more often show a pattern of startle similar to the averaged RSM of startle considering intraindividual variability.

At an individual level, results revealed evidence for a lack of a positive association of the individual RSM of the startle with both the NN model (Mean = −0.01, t(60)=-1.03, p = .31, BF_10_= 0.23) nor the AK model of valence (Mean = .034, t(60)=1.91, p =.059, BF_10_= 0.77; regression analysis: NN model: Mean = −.01, t(60)=-0.82, p = .41, BF_10_= 0.19; AK model: Mean = .034, t(60)= 2.09, p =.04, BF_10_=1.06; Figure S7D). Moreover, similar evidence between the individual RSM of startle and the AK model was found after controlling for time-based (Mean = .032, t(60)= 1.87, p = .066, BF_10_=0.71), valence-based categorical (Mean = .034, t(60)= 1.96, p =.058, BF_10_= 0.83), and arousal-based categorical models (Mean = .034, t(60)= 1.93, p =.058, BF_10_= 0.79) and arousal-based dimensional model (Mean = .035, t(60)= 2.03, p = .04, BF_10_= 0.95; Figure S7I).

**Figure S7.**
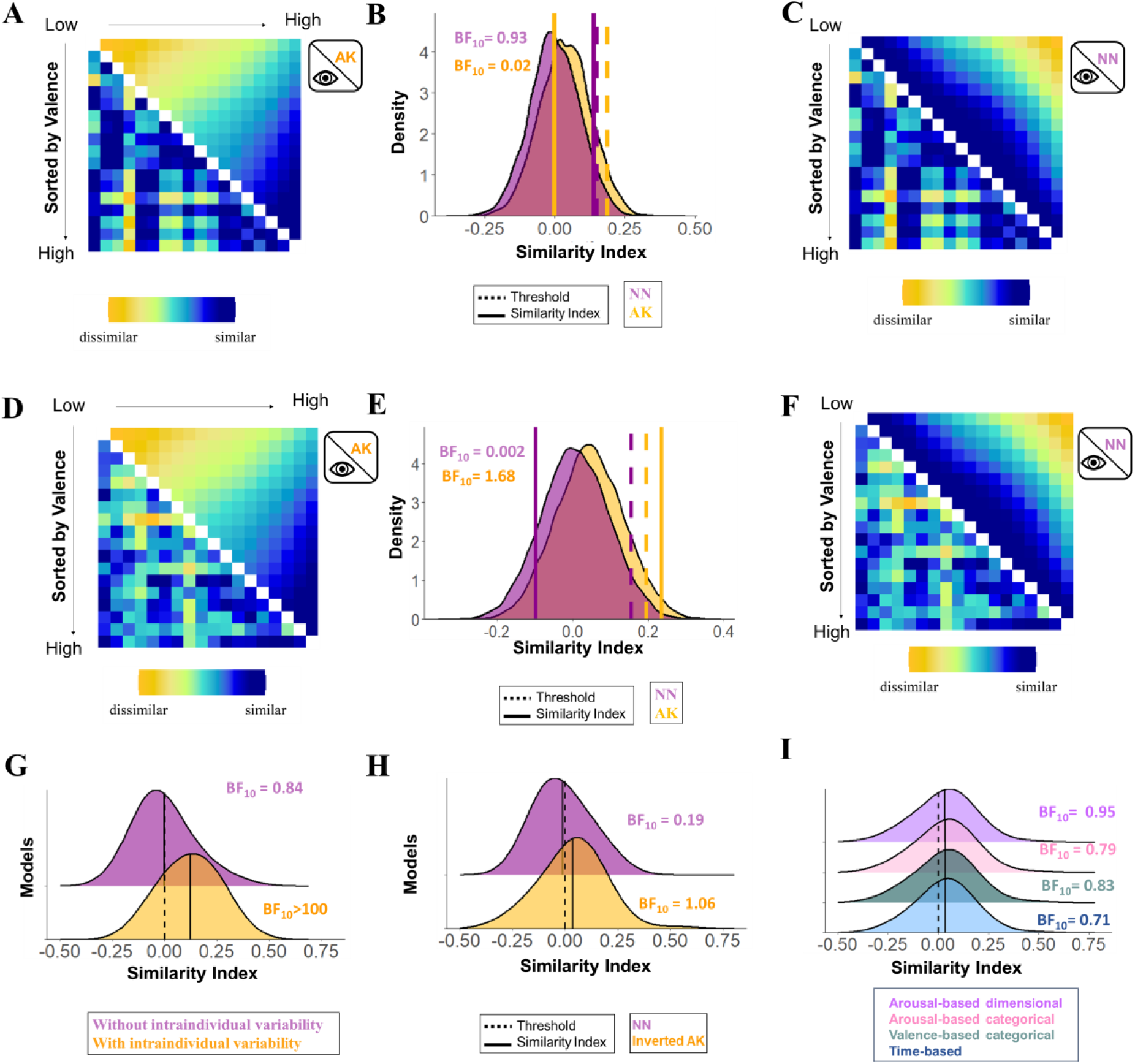
Association between the representational similarity matrices (RSMs) of subjective valence based on the Anna Karenina (AK) and Nearest Neighbours (NN) models (after controlling for the shared similarities), and the RSMs the startle eye blink response dismissing (A-C) and considering (D-F) intraindividual variability (replication sample, PSL task, N= 64). A) Averaged RSM of startle dismissing intraindividual variability (lower diagonal) and averaged RSM of subjective valence based on the AK model (upper diagonal). B) Results of the permutation test. C) Averaged RSM of startle dismissing intraindividual variability (lower diagonal) and averaged RSM of subjective valence based on the NN model (upper diagonal). D) Averaged RSM of startle considering intraindividual variability (lower diagonal) and averaged RSM of subjective valence based on the AK model (upper diagonal). E) Results of the permutation test. F) Averaged RSM of startle considering intraindividual variability (lower diagonal) and averaged RSM of subjective valence based on NN model (upper diagonal). G) Distribution of the association between individual RSMs of startle with the unique contribution of the averaged RSMs of startle disregarding (purple) and considering (yellow) intraindividual variability. H) Distribution of the association between individual RSMs of startle and unique contribution of the individual RSM of valence based on the AK and NN models. I) Distribution of the association between individual RSMs of startle and individual RSMs of subjective valence based on the AK model after controlling for other models (i.e., time-based, valence-based categorical, arousal-based categorical, valence-based dimensional models).

### 9. Representational Similarity Analysis for the association between startle and valence-based models during the Imagery task

For the Imagery task, both averaged RSMs of startle (i.e., with and without intraindividual variability) were related to each other, rho = .30, p <.001, BF_10_= 19.91, For the averaged RSM of startle dismissing intraindividual variability, moderate evidence for a lack of a positive association with the NN model of valence was found, rho = .09, p = .17, BF_10_= 0.20, and decisive evidence for lack of a positive relationship with the AK model of valence, rho = −.06, p = .81, BF_10_ <.001). When models were regressed out, and the correspondence between the model residuals and the averaged RSM of startle was tested, it was found moderate evidence for a lack of correspondence between the averaged RSM of startle dismissing intraindividual variability and the NN model of valence (rho = .1, p = .13, BF_10_= 0.30), and decisive evidence for a lack of correspondence the AK model of valence (rho =-.08, p =.84, BF_10_= 0.001 Figure S8A-C).

When the averaged RSM of startle considering intraindividual variability was compared to the models of valence, no evidence for a correspondence between the averaged RSM of startle and the NN and AK models was found e (NN model: rho = .11, p = .17, BF_10_= 0.51; AK model: rho = .19, p =.06, BF_10_= 0.41; regression analysis: NN model: rho = .08, p = .222, BF_10_= 0.42; AK model: rho = .17, p =.078, BF_10_= 0.36; Figure S8D-F).

When the averaged RSMs of startle were compared to the individual RSMs, results revealed anecdotal evidence for a lack of positive relationship between the individual RSMs of startle and the averaged RSM of startle dismissing intraindividual variability, Mean = 0.03, t(61)= 1.87, p =.06, BF_10_= 0.71, but decisive evidence for an association with the averaged RSM of startle considering intraindividual variability, Mean = 0.12, t(61)= 6.69, p <.001, BF_10_>100. The results remained when models were regressed out (averaged RSM of startle dismissing intraindividual variability: Mean = −0.007, t(61)= −0.54, p = .60, BF_10_= 0.16; averaged RSM of SCR considering intraindividual variability: Mean = 0.12, t(61)= 6.81, p <.001, BF_10_>100; Figure S8G), indicating that participants more often show a pattern of SCR similar to the averaged RSM of SCR considering intraindividual variability.

At the individual level evidence for a lack of a positive association between individual RSMs of startle and models of valence was found (NN model: Mean = .01, t(61)=1.12, p = .26, BF_10_= 0.21; AK model: Mean = .02, t(60)= 0.96, p =.39, BF_10_= 0.25; regression analysis: NN model: Mean = .01, t(61)= 1.23, p = .22, BF_10_= 0.28; AK model: Mean = .02, t(61)= 0.83, p =.40, BF_10_=0.19; Figure S7H). Moreover, no evidence for the association between the individual RSM of startle and the AK model was found after controlling for time-based (Mean = .02, t(61)= 1.00, p = .32, BF10=0.23), valence-based categorical (Mean = .02, t(61)= 1.00, p =.32, BF_10_= 0.21), arousal-based categorical models (Mean = .02, t(61)= 1.01, p =.32, BF_10_= 0.23), and arousal-based dimensional model (Mean = .02, t(61)= 0.96, p = .34, BF_10_= 0.21 Figure S8I).

**Figure S8.**
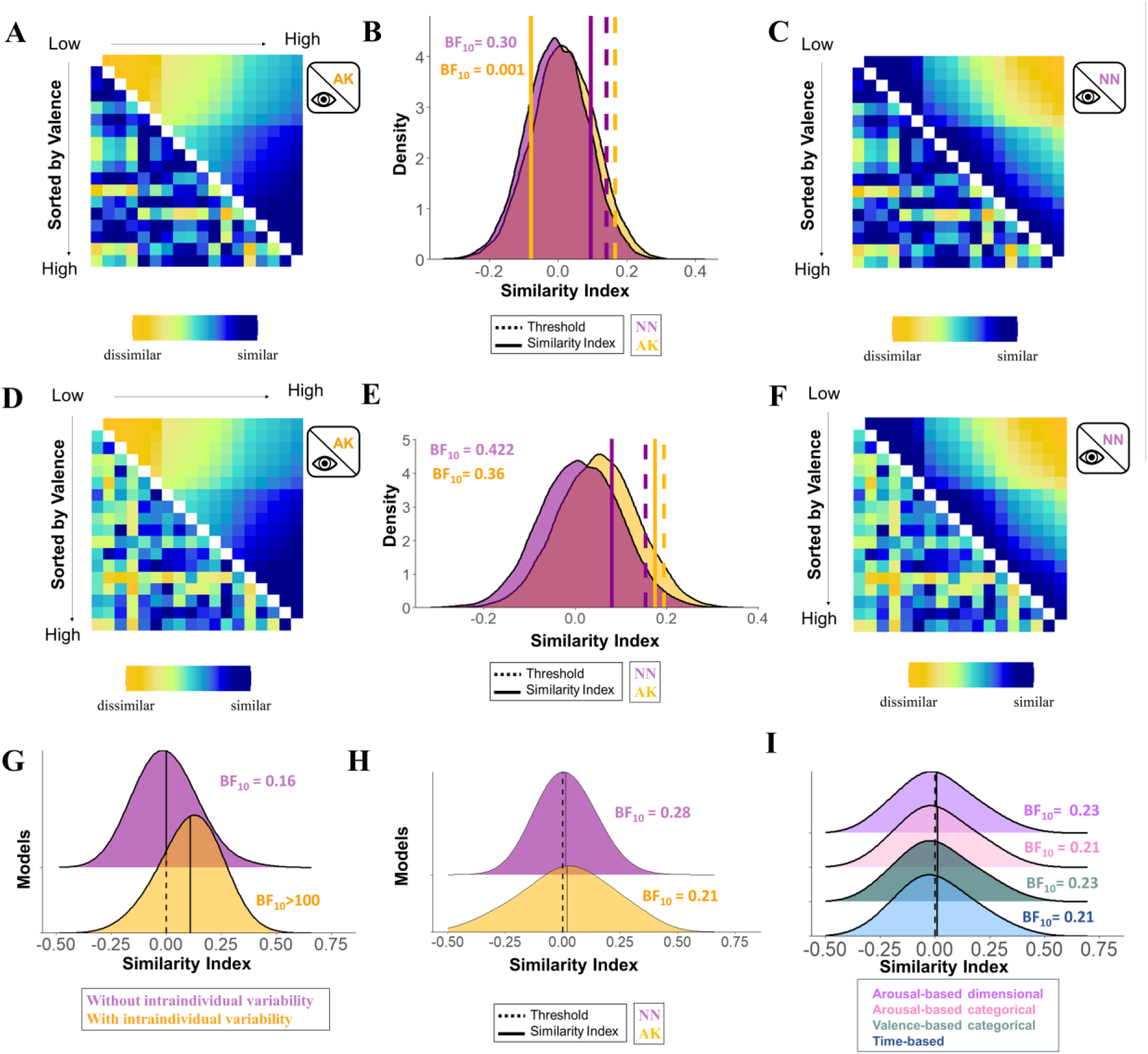
Association between the representational similarity matrices (RSMs) of subjective valence based on the Anna Karenina (AK) and Nearest Neighbours (NN) models (after controlling for the shared similarities), and the RSMs the startle eye blink response dismissing (A-C) and considering (D-F) intraindividual variability (replication sample, Imagery task, N= 64). A) Averaged RSM of startle dismissing intraindividual variability (lower diagonal) and averaged RSM of subjective valence based on the AK model (upper diagonal). B) Results of the permutation test. C) Averaged RSM of startle dismissing intraindividual variability (lower diagonal) and averaged RSM of subjective valence based on the NN model (upper diagonal). D) Averaged RSM of startle considering intraindividual variability (lower diagonal) and averaged RSM of subjective valence based on the AK model (upper diagonal). E) Results of the permutation test. F) Averaged RSM of startle considering intraindividual variability (lower diagonal) and averaged RSM of subjective valence based on NN model (upper diagonal). G) Distribution of the association between individual RSMs of startle with the unique contribution of the averaged RSMs of startle disregarding (purple) and considering (yellow) intraindividual variability. H) Distribution of the association between individual RSMs of startle and unique contribution of the individual RSM of valence based on the AK and NN models. I) Distribution of the association between individual RSMs of startle and individual RSMs of subjective valence based on the AK model after controlling for other models (i.e., time-based, valence-based categorical, arousal-based categorical, valence-based dimensional models).

## Footnotes

1 The conceptualization of the current project occurred after the two studies were conducted independently. The decision to reanalyze both studies together was based on their numerous similarities (e.g., similar physiological measures, tasks, sample demographics). However, a by-product of this *a posteriori* decision is that studies also differ in other aspects (e.g., the number of stimuli presented, the frequency of stimulus presentation, and the number of tasks used). Although some of these differences were evaluated and found to be unimportant for the results (e.g., see Suppl. S3), they should still be acknowledged.

2 We refer to affect as a basic conscious experience that can be reduced to the dimensions of valence and arousal. On the other hand, we refer to discrete emotions as complex conceptual categories that emerge out of more basic psychological operations, including but not limited to valence and arousal characteristics^16, 87^.

3 It should be noted that in the discovery sample, participants watched images twice. We decided to average the SCR across runs to have a single SCR value for each image and participant, in line with the original publication. It should, however, be noted that preliminary analysis using data from the first run led to very similar results to the ones here reported.

