## Supplementary Material for "Physiological Harmony or Discord? Unveiling the Correspondence Between Subjective Arousal, Valence and Physiological Responses"

1. Linear mixed models of the effects of Category on SCR and Startle.
2. Comparison between time-series- and amplitude-based RSMs of SCR.
3. Consistency of the time-series- and amplitude-based RSM of SCR across tasks and samples.
4. Representational Similarity Analysis for the association between SCR and arousal-based models during the PPV task in the replication sample
5. Representational Similarity Analysis for the association between SCR and arousal-based models during the PSL task in the replication sample.
6. Representational Similarity Analysis for the association between SCR and arousal-based models during the Imagery task in the replication sample.
7. Representational Similarity Analysis for the association between startle and valence-based models during the PPV task in the replication sample.
8. Representational Similarity Analysis for the association between startle and valence-based models during the PSL task in the replication sample.
9. Representational Similarity Analysis for the association between startle and valence-based models during the Imagery task in the replication sample.
1. Linear mixed models of the effects of Category on SCR and Startle

To ensure that the effects of the current data sets are in line with previous studies, showing affect-specific modulations in SCR and startle responses^1–3^, we carried out Linear Mixed Effects Models, using the lme4-package^4^. The categorical independent variable was valence category (i.e., pleasant, unpleasant or neutral; neutral was used as reference), whereas the dependent variable was z-transformed SCR and EMG amplitudes (discovery sample) or log-transformed mean phasic driver (replication sample) as well as t-transformed startle magnitudes (in both samples).

*Discovery Sample*

A significant main effect of picture valence emerged in both startle and SCR. For startle, the effect was significant (*F*(2, 20,077) = 268.48, *p* < .001), with a pattern of negative valence > neutral valence > positive valence (all pairwise comparisons *p* < .001). Similarly, for SCR, a significant effect was observed (*F*(2, 37,505) = 65.78, *p* < .001), with the valence gradient of negative > positive > neutral (all pairwise comparisons *p* < .002). startle responses aligned with the expected valence gradient, while SCR responses followed the expected arousal gradient.

Figure S1 depicts the RSM of time-series- (below the diagonal) and amplitude-based (above the diagonal) RSMs as well as the permutation test, comparing their correspondence in each of the three tasks.

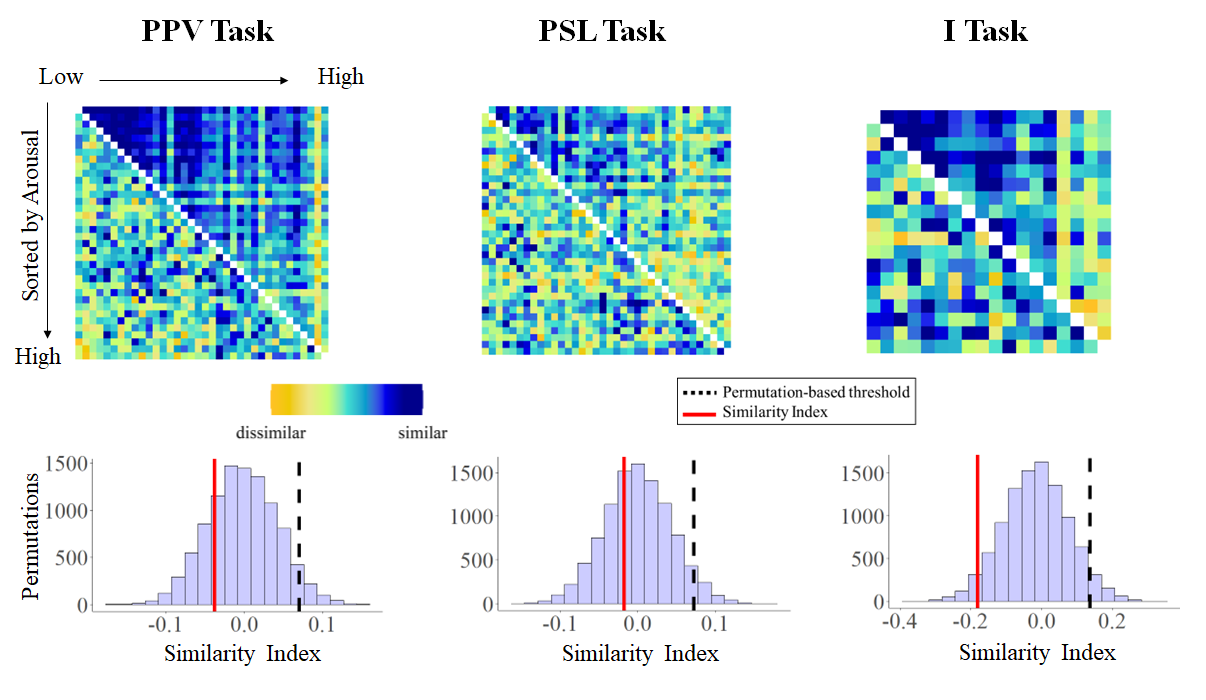

Figure S1. Results of the permutation tests comparing the RSMs derived from time-series and amplitudes of the SCR. In the upper part, the RSMs for both time-series (below the diagonal) and amplitudes (above the diagonal) are depicted for each task, PPV (right), PSL (middle), and Imagery tasks (left). In the lower part, the results of the permutation tests are depicted whereby the distribution of the permutation tests and the permutation-based threshold (black dotted line) as well as the similarity between both RSMs (red solid line) are represented. Results revealed no significant correspondence between RSMs across tasks, indicating that they are influenced by different factors.

3. Consistency of the time-series- and amplitude-based RSM of SCR across tasks and samples.

Because no correspondence was found between time-series- and amplitude-based RSMs of SCR, in the following step, we examined the consistency of both parameters by comparing RSMs from the PPV and PSL tasks. For the time-series RSMs, results revealed no similarity between RSMs (rho= .01, threshold = .11, p = .84; BF_10_ = .002). For the amplitude RSMs, however, a significant correspondence was observed, rho = .18, threshold: rho = .11, p <.001, BF_10_ = 6.69). The amplitude-based RMSs not only showed consistency between tasks within the same sample, but also significant correspondence when comparing the RSMs of the PPV and PSL task with the PPV task of the discovery sample (PPV discovery sample and PPV replication sample: rho = .39, threshold: rho = .11, p <.001, BF_10_ > 100; PPV discovery sample and PSL replication sample: rho = .26, threshold: rho = .10, p <.001, BF_10_ > 100).

Figure S2 depicts the RSM of the amplitude-based across tasks and samples as well as the permutation test, comparing their correspondence.

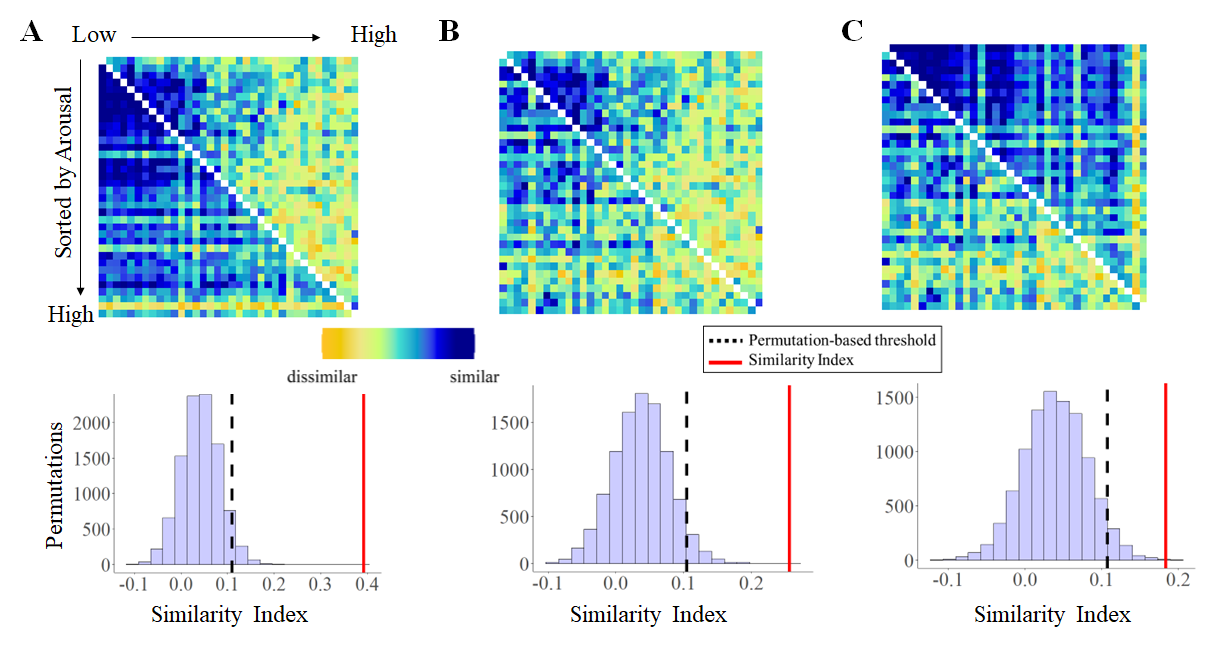

Figure S2. Results of the permutation tests comparing the RSMs derived from amplitudes of the SCR across tasks and samples. A) In the upper part, the RSMs for the SCR from the PPV of the discovery sample (above the diagonal) and from the PPV of the replication sample (below the diagonal). B) In the upper part, the RSMs for the SCR from the PPV of the discovery sample (above the diagonal) and from the PSL of the replication sample (below the diagonal). C) In the upper part, the RSMs for the SCR from the PPV (above the diagonal) and from the PSL of the replication sample (below the diagonal). In all sections (A, B, and C), the lower part depicts the results of the permutation tests whereby the distribution of the permutation tests and the permutation-based threshold (black dotted line) as well as the similarity between both RSMs (red solid line) are represented. Results revealed evidence for a correspondence between RSMs across tasks, and samples, indicating that they reflect similar processes.

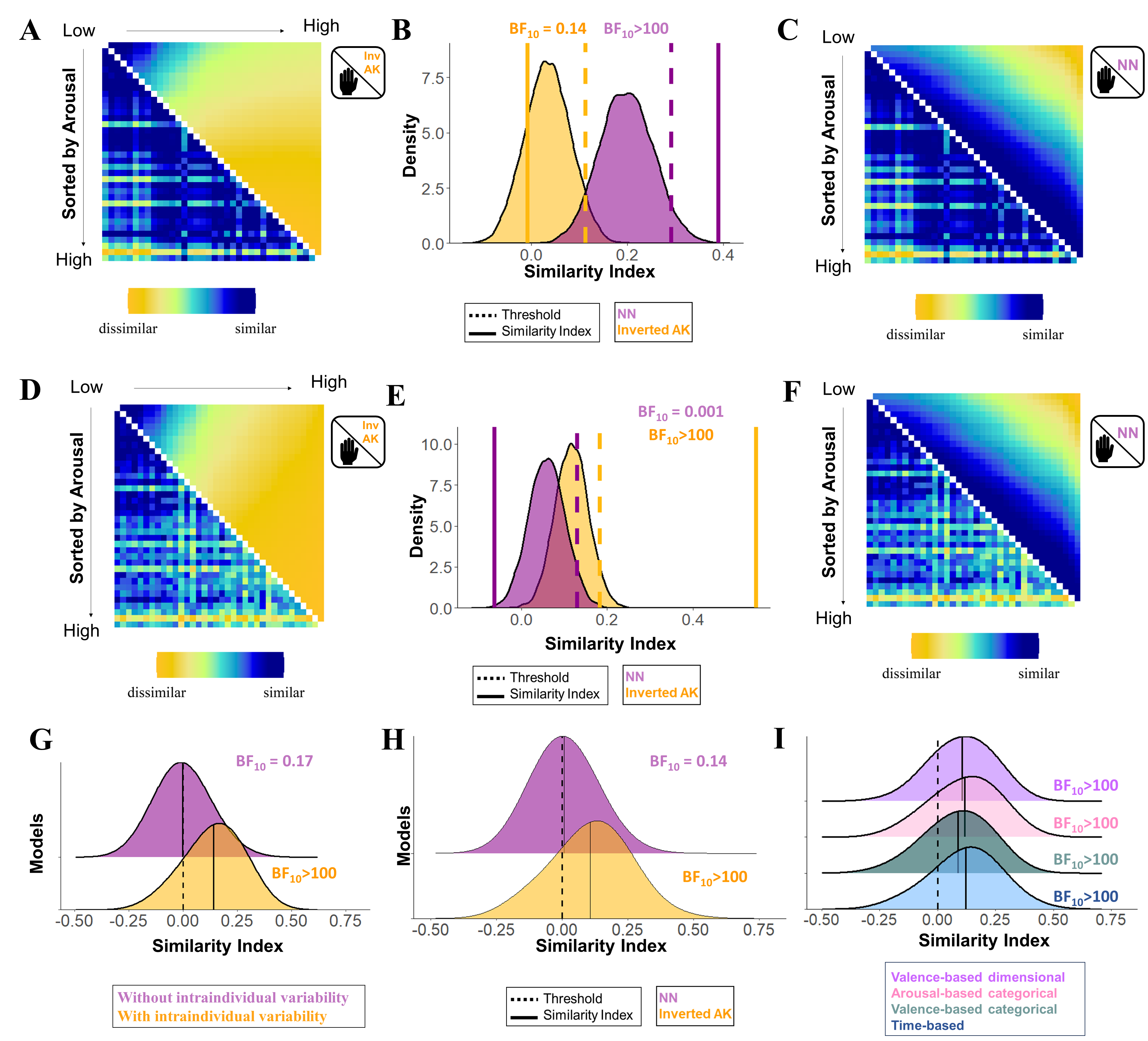

Figure S3. Association between the representational similarity matrices (RSMs) of subjective arousal based on the inverted Anna Karenina (inv AK) and Nearest Neighbors (NN) models (after controlling for the shared similarities), and the RSMs of the skin conductance response (SCR) dismissing (A-C) and considering (D-F) intraindividual variability (replication sample, N= 64, PPV task). A) Averaged RSM of SCR dismissing intraindividual variability (lower diagonal) and averaged RSM of subjective arousal based on the inverted AK model (upper diagonal). B) Results of the permutation test. C) Averaged RSM of SCR dismissing intraindividual variability (lower diagonal) and averaged RSM of subjective arousal based on the NN model (upper diagonal). D) Averaged RSM of SCR considering intraindividual variability (lower diagonal) and averaged RSM of subjective arousal based on the inverted AK model (upper diagonal). E) Results of the permutation test. F) Averaged RSM of SCR considering intraindividual variability (lower diagonal) and averaged RSM of subjective arousal based on NN model (upper diagonal). G) Distribution of the association between individual RSMs of SCR with the unique contribution of the averaged RSMs of SCR disregarding (purple) and considering (yellow) intraindividual variability. H) Distribution of the association between individual RSMs of SCR and unique contribution of the individual RSM of arousal based on the inverted AK and NN models. I) Distribution of the association between individual RSMs of SCR and individual RSMs of subjective arousal based on the inverted AK model after controlling for other models (i.e., time-based, valence-based categorical, arousal-based categorical, valence-based dimensional models).

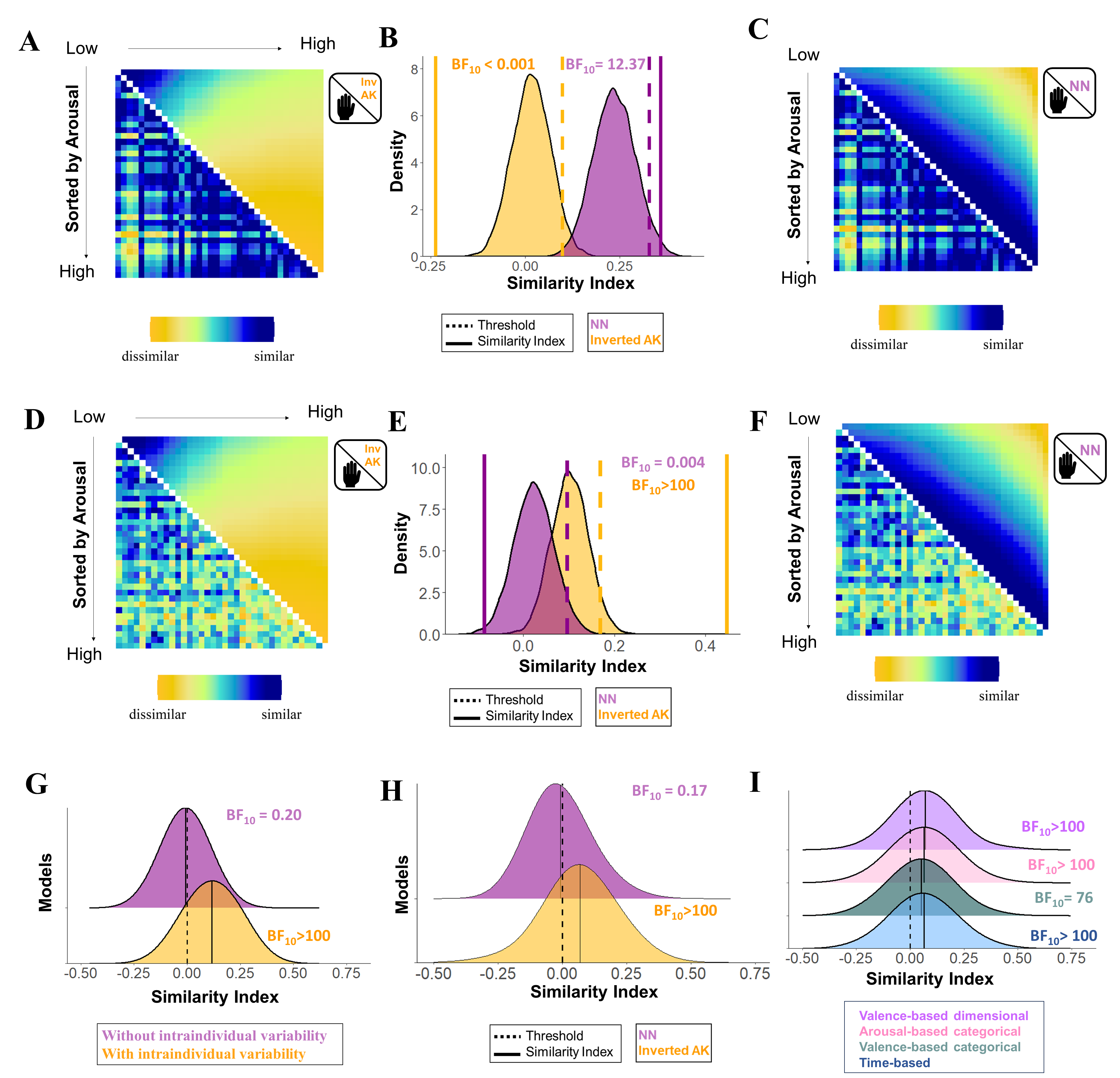
Figure S4. Association between the representational similarity matrices (RSMs) of subjective arousal based on the inverted Anna Karenina (inv AK) and Nearest Neighbors (NN) models (after controlling for the shared similarities), and the RSMs of the skin conductance response (SCR) dismissing (A-C) and considering (D-F) intraindividual variability (replication sample, N= 64, PSL task). A) Averaged RSM of SCR dismissing intraindividual variability (lower diagonal) and averaged RSM of subjective arousal based on the inverted AK model (upper diagonal). B) Results of the permutation test. C) Averaged RSM of SCR dismissing intraindividual variability (lower diagonal) and averaged RSM of subjective arousal based on the NN model (upper diagonal). D) Averaged RSM of SCR considering intraindividual variability (lower diagonal) and averaged RSM of subjective arousal based on the inverted AK model (upper diagonal). E) Results of the permutation test. F) Averaged RSM of SCR considering intraindividual variability (lower diagonal) and averaged RSM of subjective arousal based on NN model (upper diagonal). G) Distribution of the association between individual RSMs of SCR with the unique contribution of the averaged RSMs of SCR disregarding (purple) and considering (yellow) intraindividual variability. H) Distribution of the association between individual RSMs of SCR and unique contribution of the individual RSM of arousal based on the inverted AK and NN models. I) Distribution of the association between individual RSMs of SCR and individual RSMs of subjective arousal based on the inverted AK model after controlling for other models (i.e., time-based, valence-based categorical, arousal-based categorical, valence-based dimensional models).

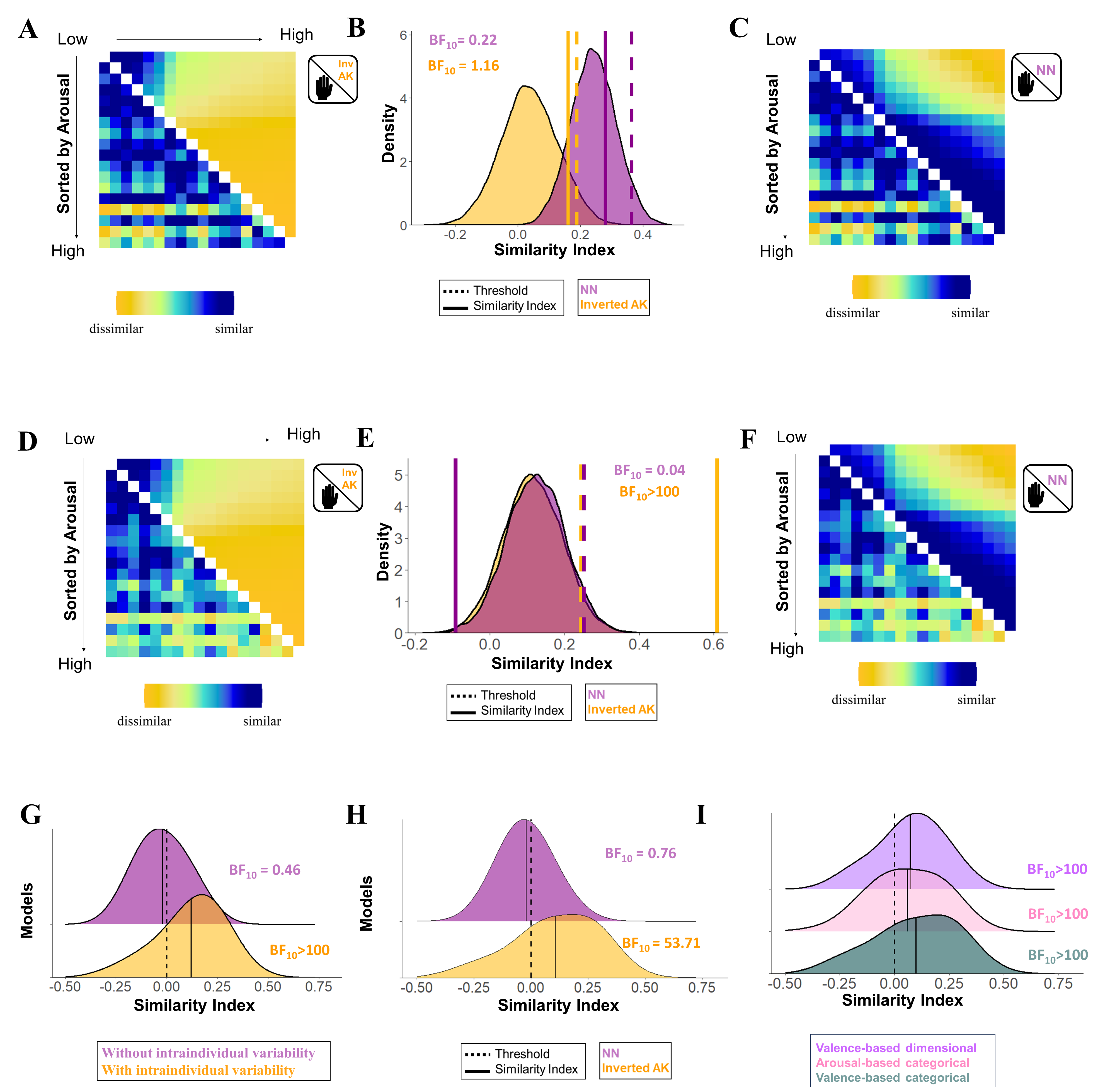
Figure S4. Association between the representational similarity matrices (RSMs) of subjective arousal based on the inverted Anna Karenina (inv AK) and Nearest Neighbours (NN) models (after controlling for the shared similarities), and the RSMs of the skin conductance response (SCR) dismissing (A-C) and considering (D-F) intraindividual variability (replication sample, Imagery task, N= 64). A) Averaged RSM of SCR dismissing intraindividual variability (lower diagonal) and averaged RSM of subjective arousal based on the inverted AK model (upper diagonal). B) Results of the permutation test. C) Averaged RSM of SCR dismissing intraindividual variability (lower diagonal) and averaged RSM of subjective arousal based on the NN model (upper diagonal). D) Averaged RSM of SCR considering intraindividual variability (lower diagonal) and averaged RSM of subjective arousal based on the inverted AK model (upper diagonal). E) Results of the permutation test. F) Averaged RSM of SCR considering intraindividual variability (lower diagonal) and averaged RSM of subjective arousal based on NN model (upper diagonal). G) Distribution of the association between individual RSMs of SCR with the unique contribution of the averaged RSMs of SCR disregarding (purple) and considering (yellow) intraindividual variability. H) Distribution of the association between individual RSMs of SCR and unique contribution of the individual RSM of arousal based on the inverted AK and NN models. I) Distribution of the association between individual RSMs of SCR and individual RSMs of subjective arousal based on the inverted AK model after controlling for other models (i.e., valence-based categorical, arousal-based categorical, valence-based dimensional models).

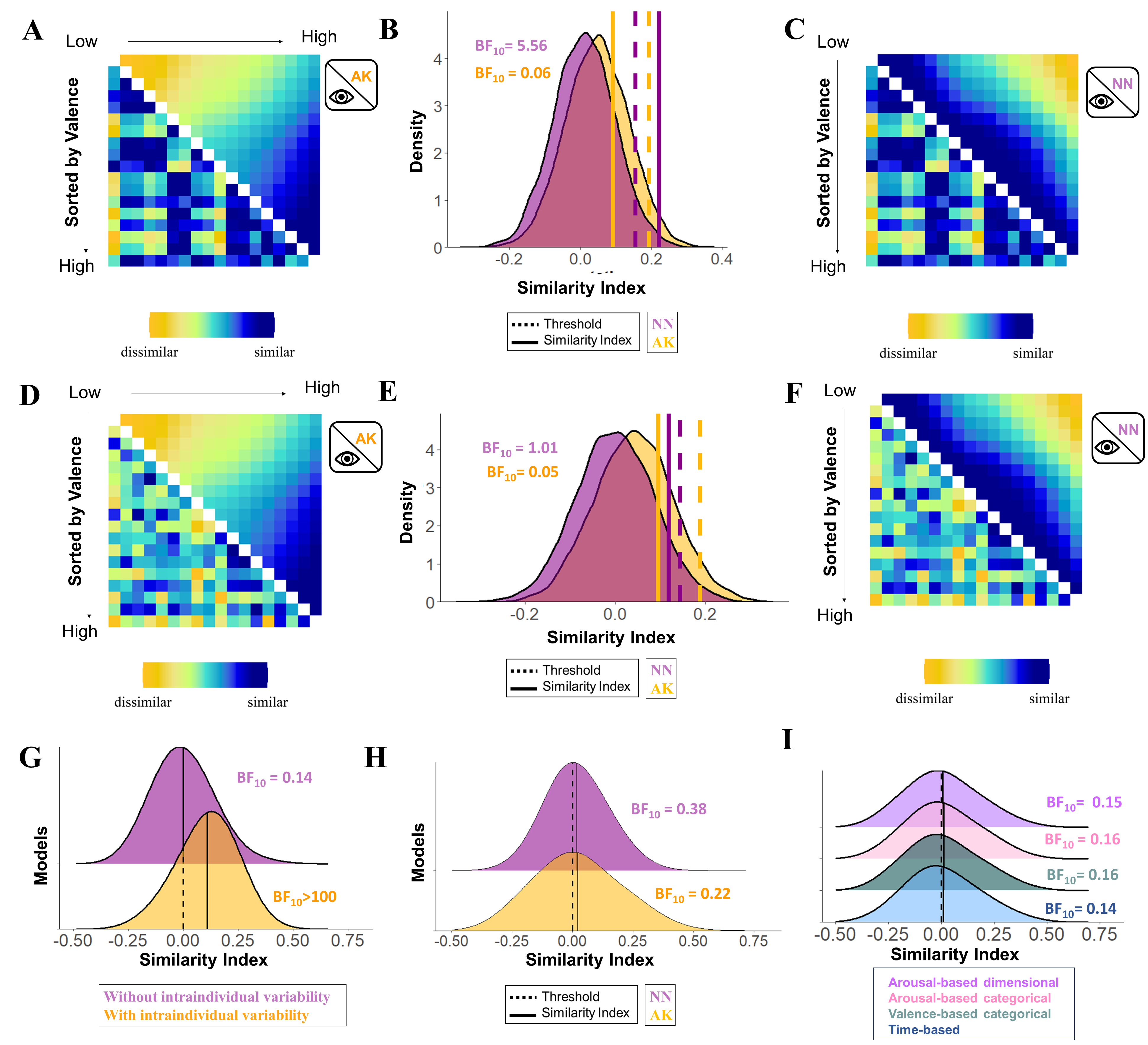

Figure S6. Association between the representational similarity matrices (RSMs) of subjective valence based on the Anna Karenina (AK) and Nearest Neighbours (NN) models (after controlling for the shared similarities), and the RSMs the startle eye blink response dismissing (A-C) and considering (D-F) intraindividual variability (replication sample, PPV task, N= 64). A) Averaged RSM of startle dismissing intraindividual variability (lower diagonal) and averaged RSM of subjective valence based on the AK model (upper diagonal). B) Results of the permutation test. C) Averaged RSM of startle dismissing intraindividual variability (lower diagonal) and averaged RSM of subjective valence based on the NN model (upper diagonal). D) Averaged RSM of startle considering intraindividual variability (lower diagonal) and averaged RSM of subjective valence based on the AK model (upper diagonal). E) Results of the permutation test. F) Averaged RSM of startle considering intraindividual variability (lower diagonal) and averaged RSM of subjective valence based on NN model (upper diagonal). G) Distribution of the association between individual RSMs of startle with the unique contribution of the averaged RSMs of startle disregarding (purple) and considering (yellow) intraindividual variability. H) Distribution of the association between individual RSMs of startle and unique contribution of the individual RSM of valence based on the AK and NN models. I) Distribution of the association between individual RSMs of startle and individual RSMs of subjective valence based on the AK model after controlling for other models (i.e., time-based, valence-based categorical, arousal-based categorical, valence-based dimensional models).

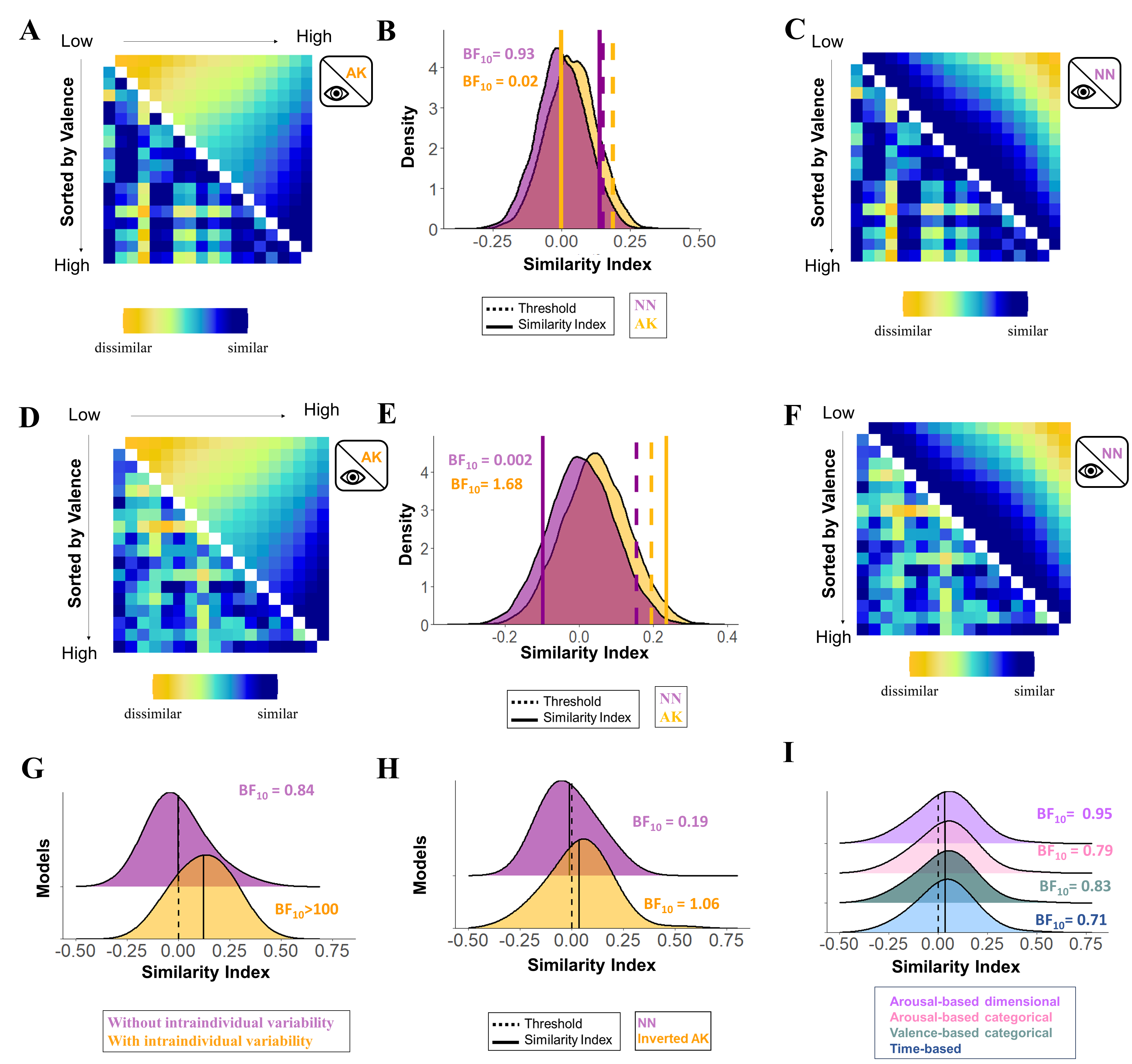

Figure S7. Association between the representational similarity matrices (RSMs) of subjective valence based on the Anna Karenina (AK) and Nearest Neighbours (NN) models (after controlling for the shared similarities), and the RSMs the startle eye blink response dismissing (A-C) and considering (D-F) intraindividual variability (replication sample, PSL task, N= 64). A) Averaged RSM of startle dismissing intraindividual variability (lower diagonal) and averaged RSM of subjective valence based on the AK model (upper diagonal). B) Results of the permutation test. C) Averaged RSM of startle dismissing intraindividual variability (lower diagonal) and averaged RSM of subjective valence based on the NN model (upper diagonal). D) Averaged RSM of startle considering intraindividual variability (lower diagonal) and averaged RSM of subjective valence based on the AK model (upper diagonal). E) Results of the permutation test. F) Averaged RSM of startle considering intraindividual variability (lower diagonal) and averaged RSM of subjective valence based on NN model (upper diagonal). G) Distribution of the association between individual RSMs of startle with the unique contribution of the averaged RSMs of startle disregarding (purple) and considering (yellow) intraindividual variability. H) Distribution of the association between individual RSMs of startle and unique contribution of the individual RSM of valence based on the AK and NN models. I) Distribution of the association between individual RSMs of startle and individual RSMs of subjective valence based on the AK model after controlling for other models (i.e., time-based, valence-based categorical, arousal-based categorical, valence-based dimensional models).

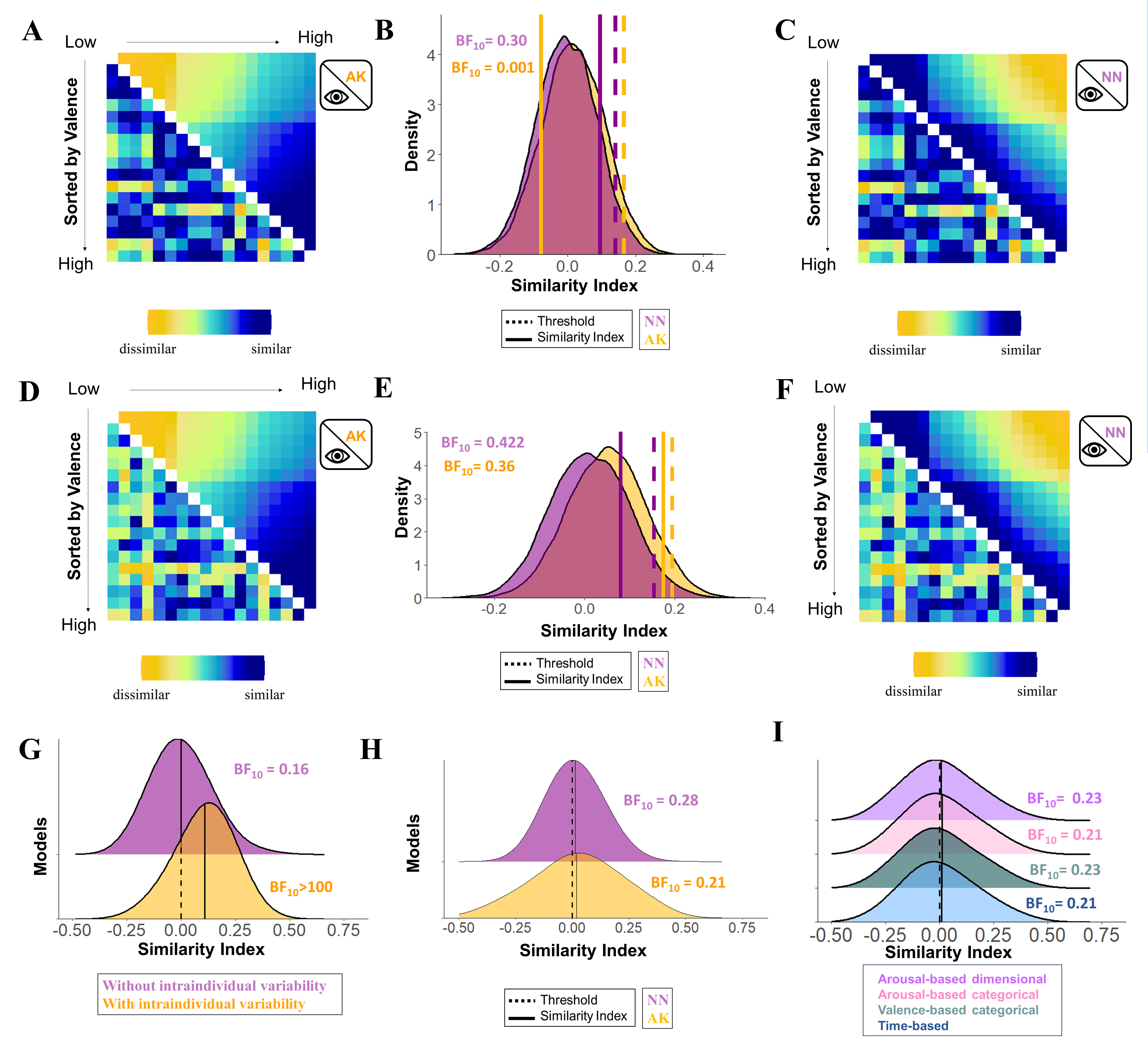

Figure S8. Association between the representational similarity matrices (RSMs) of subjective valence based on the Anna Karenina (AK) and Nearest Neighbours (NN) models (after controlling for the shared similarities), and the RSMs the startle eye blink response dismissing (A-C) and considering (D-F) intraindividual variability (replication sample, Imagery task, N= 64). A) Averaged RSM of startle dismissing intraindividual variability (lower diagonal) and averaged RSM of subjective valence based on the AK model (upper diagonal). B) Results of the permutation test. C) Averaged RSM of startle dismissing intraindividual variability (lower diagonal) and averaged RSM of subjective valence based on the NN model (upper diagonal). D) Averaged RSM of startle considering intraindividual variability (lower diagonal) and averaged RSM of subjective valence based on the AK model (upper diagonal). E) Results of the permutation test. F) Averaged RSM of startle considering intraindividual variability (lower diagonal) and averaged RSM of subjective valence based on NN model (upper diagonal). G) Distribution of the association between individual RSMs of startle with the unique contribution of the averaged RSMs of startle disregarding (purple) and considering (yellow) intraindividual variability. H) Distribution of the association between individual RSMs of startle and unique contribution of the individual RSM of valence based on the AK and NN models. I) Distribution of the association between individual RSMs of startle and individual RSMs of subjective valence based on the AK model after controlling for other models (i.e., time-based, valence-based categorical, arousal-based categorical, valence-based dimensional models).
